# Combined patterns of activity of major neuronal classes underpin a global change in brain state during spontaneous and forced walk in *Drosophila*

**DOI:** 10.1101/2022.01.17.476660

**Authors:** Sophie Aimon, Karen Y. Cheng, Julijana Gjorgjieva, Ilona C. Grunwald Kadow

**Author notes:** Max Planck Institute for Biological Cybernetics, 72076 Tübingen, Germany (Sophie Aimon).

## Abstract

Movement-correlated brain activity has been found across species and brain regions. Here, we used fast whole-brain lightfield imaging in adult *Drosophila* to investigate the relationship between walk and neuronal activity. We observed brainwide activity that tightly correlated with spontaneous bouts of walk. While imaging specific sets of excitatory, inhibitory, and neuromodulatory neurons highlighted their joint contribution, spatial heterogeneity in forward walk- and turning-induced activity allowed parsing unique responses from subregions and sometimes individual neurons. For example, previously uncharacterized serotonergic neurons were inhibited during walk. While activity onset in some areas preceded walk onset exclusively in spontaneously walking animals, spontaneous and forced walk elicited highly similar activity in most brain regions. These data suggest a major contribution of walk and walk-related sensory or proprioceptive information to brain state. We conclude that walk-related signals induce global but heterogeneous patterns of activity, allowing for local and brain-wide integration of behavioral state into other processes in the brain.

## Introduction

Growing evidence from nematodes to mammals shows that ongoing behavior affects brain activity globally (Kaplan and Zimmer, 2020; Parker et al., 2020). Using a combination of imaging and neuropixel recordings in awake, behaving mice, recent work showed that multiple dimensions of (spontaneous) behavior, including facial or body movements, are represented brainwide, allowing the integration of external or internal stimuli with the current behavioral state (Mace et al., 2018; Musall et al., 2019; Stringer et al., 2019). Brainwide imaging at single cell-resolution of calcium activity in *C. elegans* and larval zebrafish indicated that as in mammals, multiple aspects of behavior and motor activity are represented across the brain including areas thought to be dedicated to primary sensory information processing (Kato et al., 2015; Marques et al., 2020; Vladimirov et al., 2014). Importantly, such brainwide representations of motor states are seen independent of visual or olfactory inputs.

Previous studies suggested a similar situation in insects. For example, in *Drosophila melanogaster,* active flight modulates visual motion processing (Maimon et al., 2010). Similarly, visual horizontal system cells encode quantitative information about the fly’s walking behavior independently of visual input (Fujiwara et al., 2017). Beyond primary sensory brain areas, several types of dopaminergic neurons innervating the fly’s higher brain center, the mushroom body (MB), show activity highly correlated with bouts of walking (Siju et al., 2020; Zolin et al., 2021). Importantly, wholebrain imaging revealed that behavior-related activity occurred in most brain regions and was independent of visual or olfactory input (S Aimon et al., 2019; Mann et al., 2021; Schaffer et al., 2021).

These and other studies provide convincing evidence for brainwide signatures of behavior across species. However, how behavior-related information is relayed to the brain, how it spreads within neural networks and what it represents remain largely unanswered questions. Complementary to other animal models, the fly provides opportunities to study adult, global brain states associated with behavior at high temporal and spatial resolution due to its small brain size (Aimon and Grunwald Kadow, 2019). In addition, recent electron microscopy (EM) connectomics highlighted that neural networks spread across the entire brain with many pathways unrevealed by traditional single neuron manipulation experiments (Scheffer et al., 2020). Adult *Drosophila* behavior has been studied in detail for many years (Berman et al., 2016; Calhoun et al., 2019; DeAngelis et al., 2019; Geurten et al., 2014; Katsov et al., 2017; Mendes et al., 2014; Mueller et al., 2019; Tao et al., 2019; Tsai and Chou, 2019), but the resulting or underlying brainwide activity for even just roughly defined behavioral states such as resting or walking remains largely unknown.

Movement in adult *Drosophila* is thought to be controlled by descending neurons connecting the brain to the ventral nerve cord (VNC), which contains local pattern generators (Bidaye et al., 2020, 2014; Emanuel et al., 2020; von Philipsborn et al., 2011; Zacarias et al., 2018). This top-down view is challenged by other studies suggesting that behavioral control is decentralized with feedback loops involving the brain (Schilling and Cruse, 2020; Sims et al., 2019), or even that many behaviors could be locally controlled by neurons in the VNC without involving the brain (e.g., decapitated grooming (Hirsh, 1997)).

Here, we build on our previous work (Aimon et al., 2019) using fast whole brain imaging during ongoing behavior to unravel more comprehensively the spatial and temporal relationship between movement and neural network activity across multiple brain structures and neuronal subtypes. We first identify aspects of the behavior encoded by regions and neurons expressing different neuromodulators. We next characterize the timing of activation at the behavior transitions between rest and walk and find that walk-related activity in most brain regions happens at or after the transition. Nevertheless, several activity components, e.g., in the posterior slope, increase their activity before walk onset. This is consistent with the hypothesis that brain activity in most regions originates primarily from efferent or proprioceptive feedback, and that few brain regions are directly involved in top-down movement control. We test this hypothesis by comparing global brain activity during forced and spontaneous walk and find them to be similar in most brain regions with select areas preceding exclusively spontaneous walk. Our results suggest that walk triggered activity in select brain areas and neurons underly a global change in brain state thereby allowing for efficient integration of external information with ongoing behavioral actions to guide appropriate future decisions through behavioral adaptation and learning.

## Results

### Whole brain imaging reveals broad activation during walk

To image whole brain activity during spontaneous behavior, we fixed the fly’s head to a holder and opened the posterior head capsule while leaving the legs free to move (Woller et al., 2021) (Fig. 1A), and used a ball as a walkable surface (Seelig et al., 2010) (Fig. 1A). We also carried out experiments without a walking substrate to compare activity related to walk to activity related to flailing movements, where the fly moves its legs freely in the air. We expressed GCaMP pan-neuronally (i.e., *nsyb-gal4;UAS-GCaMP6s/6m/6f/7s/7f*) and imaged calcium transients as a proxy of neuronal activity in the whole brain using fast light field microscopy (LFM) as described previously in detail (Sophie Aimon et al., 2019). Briefly, a multilens array captures an image of the entire brain. This raw image is further processed to reconstruct the brain volume, corrected for movement artifacts, and aligned to a template to identify the brain regions or activity components with behavior-correlated calcium changes (Fig. 1A).

**Figure 1:**
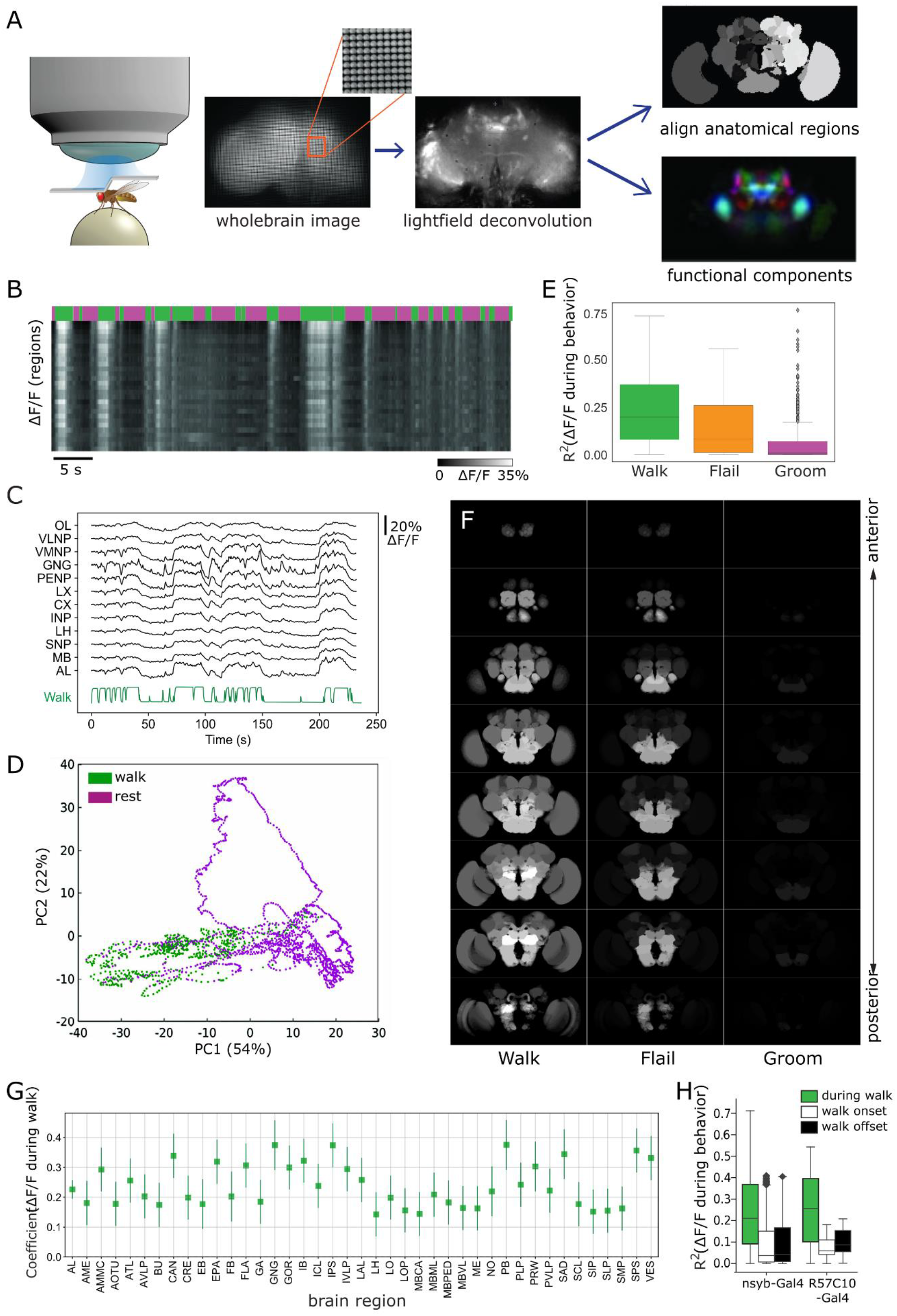
Global brain activation during walk. (**A**) Schematic overview of the preparation and analysis method. Please see methods for details. (**B**) Raster plot of the activity of regions. Top panels depict walking bouts in green and rest or grooming in magenta. Lower panel shows calcium activity elicited throughout the experiment. The brighter the higher the calcium transients. Mean forward speed: 5.6 mm/s, mean angular velocity: 0.4 radian/s (**C**) Sample traces (deltaF/F) for different brain regions relative to forward walk (green). (**D**) First two principal components from whole brain activity color-coded with behavior. (**E**) R^2^ for different behavioral conditions (all regions were pooled, but p-values are obtained after averaging regions for each fly. Walk: N=16, Flail: N=7, Groom: N=6). P-values: Walk vs. Flail 0.085, Walk vs. Groom: 0.011, Flail vs. Groom: 0.26. Center line, median; box limits, upper and lower quartiles; whiskers, 1.5x interquartile range; points, outliers (**F**) Z-stack map of R^2^ median (Walk: N (flies) =16, Flail: N=7, Groom: N=6) for regression between regional activity and walk, flail or groom. Mann-Whitney-test: walk vs. flail: 85, walk vs. groom: 85; flail vs. groom: 31 (**G**) Normalized coefficient of different regions’ activity regression with walk. All regions’ 95% CI are above zero. N=16. (**H**) R^2^-value of regression coefficients during walk, walk onset and walk offset (all regions were pooled, nsyb-Gal4: N=16, GMR57C10-Gal4: N=4). Regressors for walk onset or offset are Dirac functions convolved with the GCaMP response. Box plots show: center line, median; box limits, upper and lower quartiles; whiskers, 1.5x interquartile range; points, outliers.

In order to compare periods of rest with periods of spontaneous walk, we combined behavioral movies and z-projections of whole brain data (Movie 1). We observed a strong increase in neuronal activity across the brain during bouts of walking compared to during resting or grooming (Fig. 1B, C).

While LFM allows for very fast imaging frame rates (up to 200 Hz for the whole brain), the frame rate was limited by the signal-to-noise ratio in each individual fly and was thus adjusted for each experiment (see Methods). We found no significant difference between the global activity R^2^ for walk at different frame rates (5 – 98 Hz) for flies expressing GCaMP6f pan-neuronally (Fig. S1A). Thus, we pooled these data for the analysis below regarding global brain state.

In experiments where the fly was both walking and resting, the first principal component (PC1) of the whole brain activity data was strongly correlated to walk (Fig. 1D). Next, by aligning whole brain imaging data to an anatomical template using landmark registration, and by averaging activity in each large anatomically defined region (see Methods), we also found that the global activity of regions correlated with walk (Fig. 1B and C). Importantly, neuronal activation followed movement bouts at high temporal precision (limited by the temporal resolution of the calcium reporter) inconsistent with activity generated by general arousal, which we would not expect to precisely follow such bouts (Fig. 1B and C). However, the calcium responses were still a little slower than some of the fast changes in walking, even after convolution of walking with an impulse response, which suggests that faster calcium probes could capture finer dynamics in further studies (Fig. 1C).

We next mapped the differences in activity during walk, flail, or groom to a brain template with finer brain regions to understand their spatial organization. While we still observed an increase in activity during bouts of flailing, the brain was not as consistently globally activated as during walking (Fig. 1E and F). Strikingly, unlike grooming, which resulted in local activation of ventral brain areas consistent with (Hampel et al., 2015), all brain regions were significantly correlated with walking (Fig. 1E and F). Indeed, using walking behavior we were able to explain ~ 20 % of all variances observed in the experiments (R^2^ median = 0.194, Fig. 1E). We used a linear model to analyze how the correlation of brain activity with walking (normalized coefficient and the R^2^) depended on the brain region. To ensure our results were not biased by heterogeneous expression in the pan-neuronal transgenic driver line, we used two different pan-neuronal lines (*nsyb-Gal4* and *GMR57C10-Gal4*). The linear model (Table 1) showed that there was no significant effect of the Gal4-driver. We found a small but significant effect of the GCaMP version used, however, and thus kept using the model to take this effect into account for the rest of the study. Using this model, we quantified the effect of the distinct brain regions on both the normalized coefficient and the R^2^ of the regression with a behavior regressor (Fig. 1G, S1C-G,J,K). While all brain areas were significantly activated during walk (Fig. 1G, S1C), only the region of the gnathal ganglion (GNG) showed strong activation during grooming (Fig. S1E,G). Flailing represents an intermediate state with many brain regions activated but less consistently as compared to walk (Fig. 1E, S1D,F). Moreover, the global activity correlated primarily with walk and not with the start or end of walk (Fig. 1H).

To test whether part of the observed global brain activity could be due to visual input coupled with the behavior, e.g., by the fly seeing the optic flow from the ball (Borst et al., 2020; Suver et al., 2016), or unexpectedly fixed reflections from the environment (Creamer et al., 2018), we performed the same experiments but covered the fly’s eyes with black nail polish to prevent outside light from activating its photoreceptor neurons. With such strongly limited visual input, we still observed a very similar, global activity pattern indicating that visual input is a minor contributor to the observed wide brain activity (Fig. S1B). This also suggested that the global increase in activity during walk is not due to a mismatch between an actual and a predicted visual stimulus. In addition, given that air-supported balls can show erratic movements due to air turbulences that could cue the fly to change its behavior (DeAngelis et al., 2019; Sen et al., 2019), or impose constraints on the animal’s posture, we performed parallel experiments with an unsupported styrofoam ball held by the fly (see Methods). The comparison of the two datasets revealed a significant difference in R^2^ for global brain activity (Fig. S1H), indicating that the type of walking substrate influenced brain activation. Given that the activity distribution across brain regions did not significantly differ between the two substrate types at this level of analysis (Fig. S1I), we combined the datasets for most analysis below.

Together these data confirm that walking behavior induces a change in global brain activity (Sophie Aimon et al., 2019) with most brain regions showing highly temporally correlated activity during bouts of spontaneous walk. They further suggest that most global activity occurs during walk and not at or before its start or end.

### Both inhibitory and excitatory neurons are recruited during walk

We next asked what major neuron types underly this global change. Previous work indicated that 37% of all descending neurons (DN) to the ventral nerve cord (VNC) are GABAergic and 38% cholinergic, while 6% express the neurotransmitter glutamate (Hsu and Bhandawat, 2016). This suggests that both excitatory and inhibitory neurons influence behavior and could underly behavior-related brain states. It is also conceivable that broad brain activation arises due to a global disinhibition during periods of walking (Benjamin, 2010), in which case we would see a decrease of activity in inhibitory neurons.

To address these questions, we expressed GCaMP6m exclusively in GABAergic inhibitory neurons using the GAD1-Gal4 driver. We also did the same with a driver for glutamatergic neurons (*Vglut-gal4;UAS-GCaMP6m*) as glutamate is thought to be inhibitory in the fly. In addition, we analyzed activation patterns during behavior in excitatory neurons by imaging from all cholinergic neurons in the fly brain (*Cha-gal4;UAS-GCaMP6m or UAS-GCaMP6f*). Using the same approach and analysis as described above, we detected an increase in global brain activation, which again correlated highly with walking bouts, for excitatory and inhibitory types of neurons (Fig. 2A).

**Figure 2:**
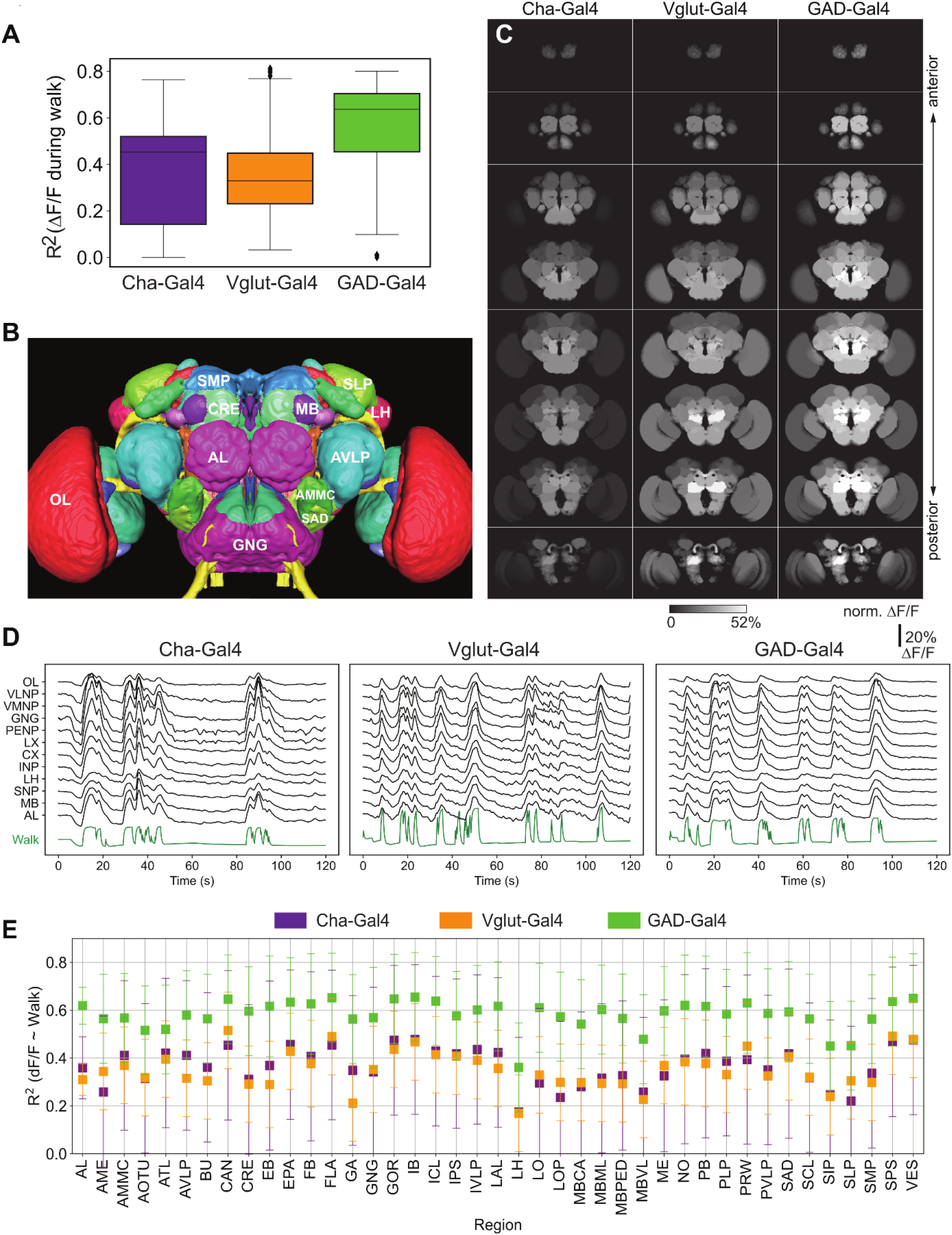
Activity of neurons releasing the three major neurotransmitters, glutamate, GABA and acetylcholine during walk. (**A**) R^2^ for regression of activity with walk in different conditions (all regions were pooled). No pairwise comparison is significantly different. Mann-Whitney-test: Cha vs. GAD: 4; Cha vs. Vglut: 15; GAD vs. Vglut: 22 (**B**) Overview of major brain regions as defined in (Ito et al., 2014). (**C**) Activity maps (regression coefficient) of functional regions activated during walk for Cha-Gal4, GAD-Gal4 and Vglut-Gal4 expressing neurons. (**D**) Sample traces (deltaF/F) for different brain regions relative to forward walk (green). (**E**) R^2^ of regression of fluorescence vs. walk for Cha-Gal4, GAD-Gal4 and Vglut-Gal4 expressing excitatory neurons. N=5 flies for each individual genotype. Box plots show: center line, median; box limits, upper and lower quartiles; whiskers, 1.5x interquartile range; points, outliers.

As for the pan-neuronal data, we mapped this activity to all brain regions to create a functional map of activity during walk (Fig. 2B,C). Among other regions, we found that the GNG, the anterior ventral lateral protocerebrum (AVLP) and the antennal mechanosensory and motor center (AMMC), known to be important for leg coordination and to be highly innervated by ascending neurons (AN) from the VNC (Chen et al., 2022; Emanuel et al., 2020; Tsubouchi et al., 2017), responded to walk for all genotypes (Fig. 2B-D). Again, brain activity closely followed individual walk bouts (Fig. 2D). Using a linear regression model, we determined that similar to the pan-neuronal data, excitatory neurons labeled by Cha-Gal4 were activated in all brain regions (Fig. 2E, S2). Brain regions such as the protocerebral bridge (PB), the cantle (CAN), the vest (VES) and the superior posterior slope (SPS) were correlated especially strongly in all three types of neurons (Fig. S2A-C).

Together these data show that inhibitory neurons as well as excitatory neurons are activated in most brain regions during walk. Moreover, these data suggest that the observed activity in pan-neuronal imaging is not mediated by a global disinhibition in the brain, but rather a result of a combined activation of different types of neurons, including excitatory and inhibitory neurons.

### Aminergic neuron activity is strongly correlated with behavior

Monoamine-releasing neuromodulatory neurons, i.e., dopamine, serotonin and octopamine (the fly’s equivalent to mammalian noradrenaline) are important to adjust behavior (van Damme et al., 2021). Among the DNs, these neuromodulatory neurons make up ~4% of all neurons (41). However, the role of these neuromodulators during movement remains to be fully elucidated. Previous data showed that both walking and flailing, but not grooming or resting, were related to an increase in dopaminergic neuron activity (DAN) in some mushroom body compartments (Siju et al., 2020; Zolin et al., 2021), and many other regions across the brain (Sophie Aimon et al., 2019). We combined TH-Gal4 with DDC-Gal4 or GMR58E04-Gal4 to cover all dopaminergic neuron types (Cohn et al., 2015; Siju et al., 2020) and used Tdc2-Gal4 for octopamine and Trh-Gal4 for serotonin.

While we observed an increase in activity for all neuromodulatory neuron types, serotonergic neurons were significantly less globally activated than dopaminergic and octopaminergic neurons during spontaneous bouts of walk (Fig. 3A and D, Movie 2-4). We also mapped the activity to brain regions (Fig. 3B). Contrary to results shown above with pan-neuronal and broad neurotransmitter lines, maps were distinctly patterned for the individual aminergic lines, as revealed by overlaid activity maps (Fig. 3B,C).

**Figure 3:**
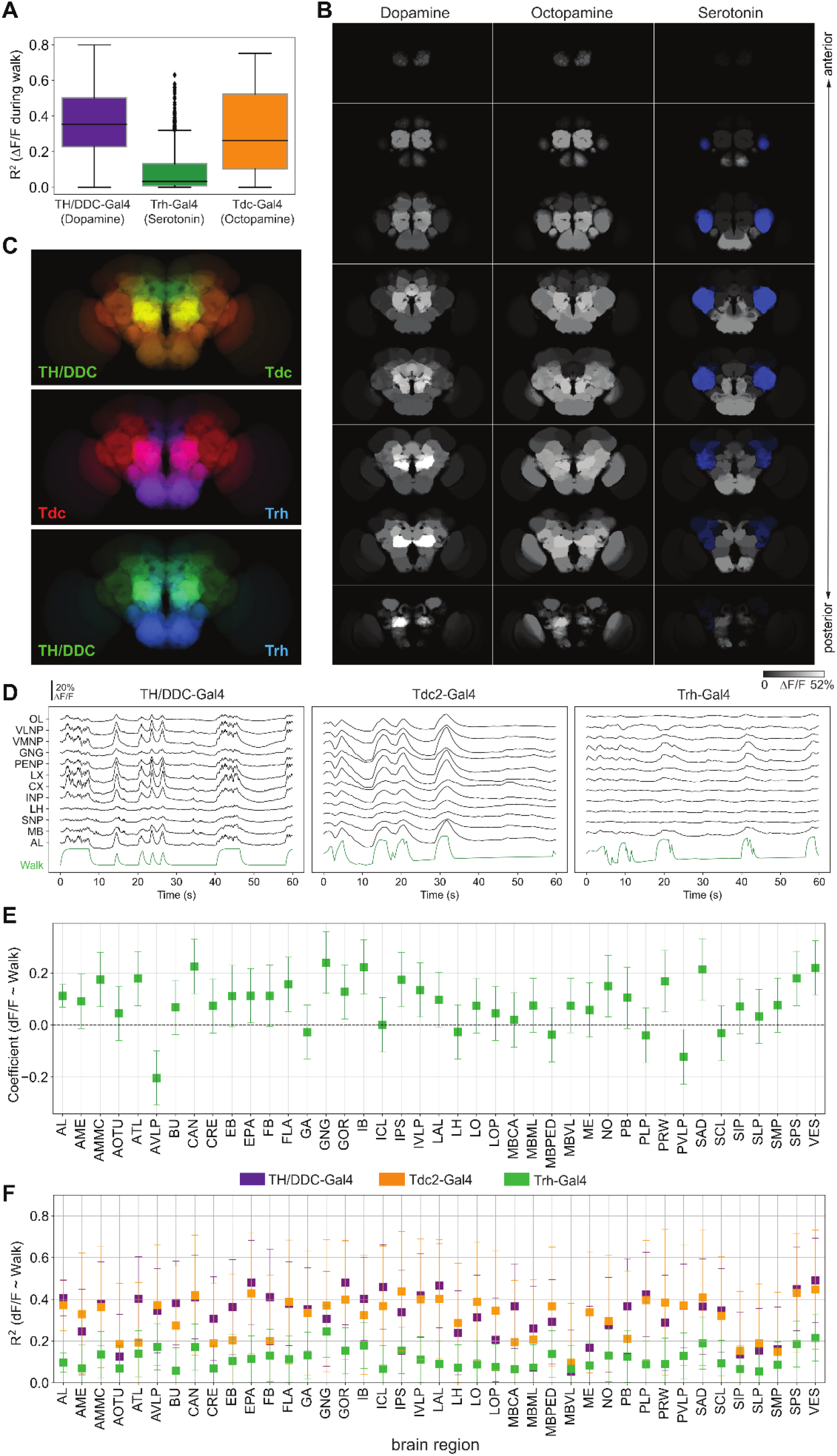
Neuromodulatory neurons are strongly and differentially activated during walk. (**A**) Coefficient of determination for behavior regression at different conditions (all regions were pooled). Mann-Whitney-test; P-values: TH vs. Trh: 47, 0.032, TH vs. TDC: 35, ns, Trh vs. TDC: 5, 0.040 (**B**) Activity maps (regression coefficient) of functional regions activated during walk for TH-gal4 and DDC-gal4 or GMR58E04-Gal4 (dopaminergic neurons), TDC2-gal4 (octopaminergic neurons) and Trh-Gal4 (serotoninergic neurons) expressing neurons. Blue indicates inhibition. (**C**) Overlay of activity maps of two neuromodulators in each panel. (**D**) Sample traces (deltaF/F) for different brain regions relative to forward walk (green). (**E**) Coefficient during walk for different brain regions for Trh-Gal4 expressing serotonergic neurons. All regions are significantly correlated. (**F**) R^2^ of regression of fluorescence vs. walk for TH/DDC(58E02)-Gal4, Tdc2-Gal4 and Trh-Gal4 expressing excitatory neurons. TH/DDC-Gal4: N=9, Tdc2-Gal4: N=7, Trh-Gal4: N=6 flies. Box plots show: center line, median; box limits, upper and lower quartiles; whiskers, 1.5x interquartile range; points, outliers.

Most regions were consistently activated by neurons expressing GCaMP under the control of TH/DDC-Gal4 (Fig. 3B-D, S3A). Importantly, besides the dopaminergic neurons that were already implicated in prior work on specific regions (Berry et al., 2015; Cohn et al., 2015; Siju et al., 2020), whole brain imaging suggested that many additional DANs were activated during or due to walk (Fig. 3A-D, (Aimon et al., 2019)). Octopaminergic neuron activity (imaged with Tdc2-gal4;UAS-GCaMP6s) was also highly correlated with walking (Fig. 3A,B,D). The activation was particularly high for neurons located or projecting into the ventromedial neuropils (VMNP, Fig. 3F and S3B).

Using whole brain imaging of serotonergic neuron activity (Trh-gal4;UAS-GCaMP6m,f or s), we found that the activity of several regions was highly correlated with walking (Fig. 3B,E,F). Surprisingly though, the strongest correlation was negative and mapped to the anterior ventrolateral protocerebrum (AVLP) region (Fig. 3B (blue area), 3E,F). This suggested that while some populations of serotonergic neurons were activated, some subsets, in particular in the AVLP, were inhibited during walk (Fig. 3E). Interestingly, the AVLP contains several descending neurons projecting toward the VNC (Namiki et al., 2022, 2018).

Together, these data show that walking activates, and in some cases inhibits, all main types of aminergic neuromodulatory neurons in various but distinct brain regions. Thus, the observed pan-neuronal global activity during walk is a result of the combined patterns of activation of excitatory, inhibitory, and neuromodulatory neuronal responses to walk.

### Whole brain activity data identifies specific brain regions or even neurons

Global brain activity can be used to generate functional maps of the brain (i.e., fMRI (Beckmann and Smith, 2004)). These functional maps not only show which brain regions are activated but provide information regarding which brain regions might be functionally connected and/or involved in similar aspects of a task due to their highly correlated activity (i.e., functional connectome). In the case of the fly, highly detailed anatomical maps (Chiang et al., 2011; Ito et al., 2014) allow to match the shape of these functional map to subregion and in some cases to single neuron types (Table 2).

To this end, we extracted functional components using PCA followed by ICA to unmix the PCA maps (see Methods) (Fig. 4A, S4A-C). In essence, ICA unmixes the PCA values in an effort to spatially separate them. We grouped smaller functional components within a larger brain region (e.g., different antennal and protocerebral bridge glomeruli), if the precision of the alignment of the template did not allow a clear assignment of individual regions. Interestingly, almost all functional components derived from recorded neuronal activity matched anatomical structures without further subdivisions or blurring of anatomical boundaries. This suggested that the activity within brain regions was more homogenous as compared to the activity in neighboring regions. In one exception, our functional data separated a larger region in the lateral neuropil, the AVLP, into smaller subregions (Fig. S4A,B, orange box) suggesting that subregions of the AVLP had different activity signatures. Of note, some components also identified more than one brain region (i.e., PENP-SLP) indicating a strong functional, i.e. synaptic, connection between these regions.

**Figure 4:**
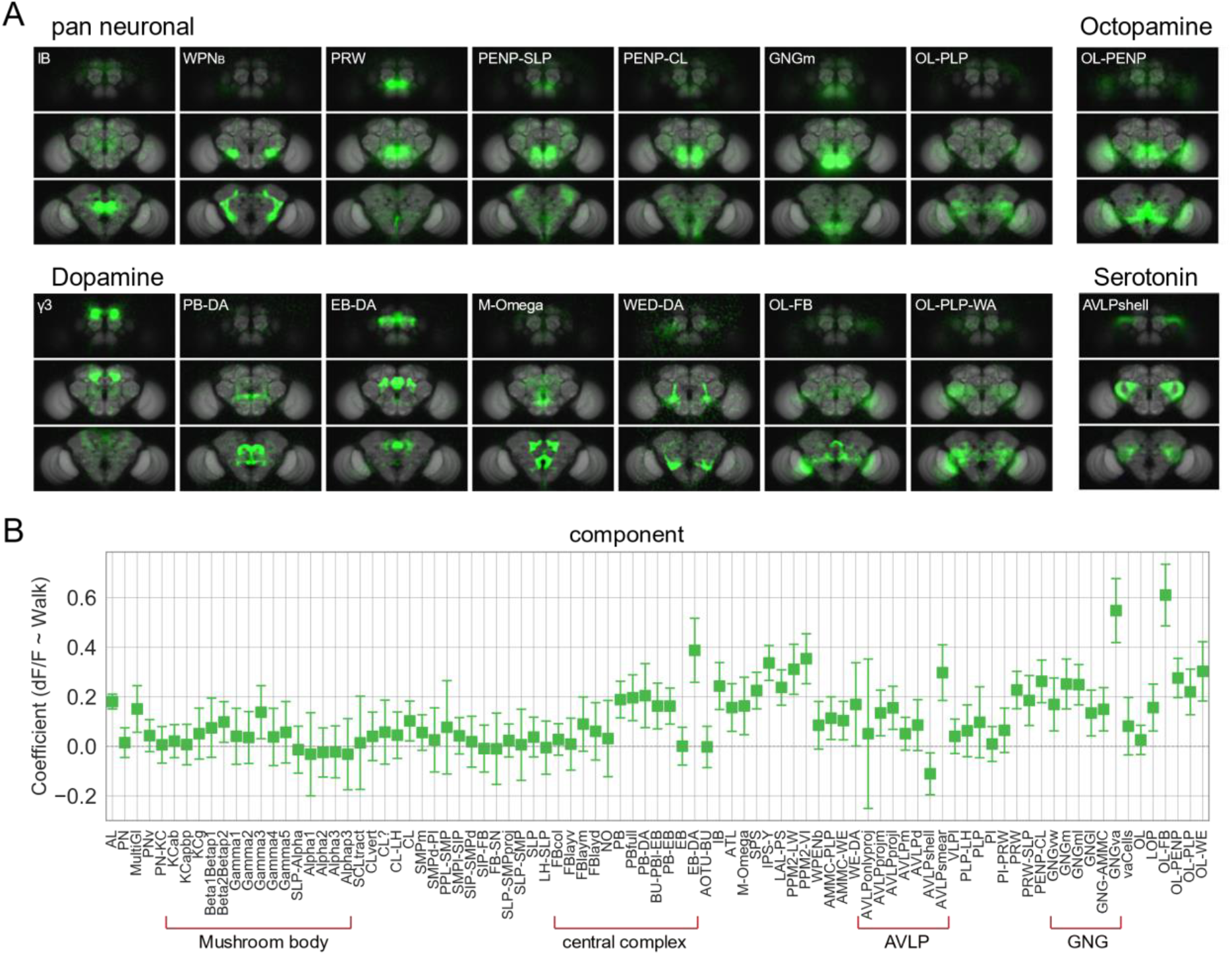
Whole brain analysis pinpoints specific subregions responding to walk. (**A**) Images of example components that are significantly correlated with walk (Coefficient or R^2^ 95% CI above zero). Upper left: components derived from imaging with a pan-neuronal driver. Upper right: components derived with Tdc2-Gal4. Lower left: components derived with TH/DDC or GMR58E02-Gal4. Lower right: components derived with Trh-Gal4. (**B**) Correlation coefficient for components activity vs walk. N=58 flies of different genotypes, see table in methods for details.

Among the components correlating with spontaneous walk, the PENP-SLP (periesophageal neuropils and superior lateral protocerebrum) and PENP-CL (periesophageal neuropils and clamp) components might contribute to relaying the information that the fly walks from the ventral neuropils (e.g., GNG) to higher areas or vice versa. The WPNb (bilateral wedge projection neuron) component (likely generated by the WPNb neuron (Coates et al., 2020)) showed weaker correlation but still had a R^2^-value significantly above zero and could thus also relay the activity between lower and higher brain regions.

As some of the data used to extract maps were from flies expressing GCaMP specifically in inhibitory, excitatory and neuromodulatory neurons (Fig. 4A), components could be matched to neurons of the same types (Table 2). For dopaminergic neurons, walk-correlated neuronal activity was found, for instance, for components in and around the MB (i.e. γ3-compartment as previously observed; (Cohn et al., 2015; Siju et al., 2020; Zolin et al., 2021)), in the central complex (protocerebral bridge (PB-DA) and ellipsoid body (EB-DA (Kong et al., 2010))), but also in ventral neuropil such as the wedge (WE-DA (Liu et al., 2017)) and neuropil connecting central regions to the optic lobes (Fig. 4A). For octopamine, one particular component connecting the optic lobe and the periesophageal neuropil was strongly correlated with walk, the OL-PENP, while for serotonin the highest R^2^-value was detected for the AVLP and components within (Fig. 4A). Most components with significant R^2^-values were positively correlated with walk with one exception (Fig. 4B,C): a specific component of the AVLP, a subregion we named the AVLPshell due to its shape, was negatively correlated with walk, i.e., was inhibited during walk (Fig. 4B, S4A-C). As this component was mostly detected in the experiment using Trh-Gal4, serotonergic neurons were likely responsible for the decrease in activity during walk (see also Fig. 3).

Although most component were correlated with walk, other had an R^2^ indistinguishable from zero thus likely representing ongoing activity unrelated to walk (Schaffer et al., 2021).

Several of these components were previously implicated in walking or its modulation, but many others were not (Table 2) indicating that whole brain imaging during behavior in combination with our analysis pipeline provides a powerful method to pinpoint new brain regions and neurons involved in the control of, or response to, walking behavior. For most components, we were able to propose specific neurons based on anatomical resemblance (Table 2). Some of these neurons can be labeled by Gal4 transgenic lines for future functional manipulation. Furthermore, the identification of functional components reveals functional similarity and connectivity of anatomically connected brain regions in spontaneous walk.

### Turning activates specific brain regions and neurons

Some functional components activated during walk showed an activity that appeared to be mirrored by the other half of the brain (Fig. 5A). We next asked whether these components had differential activity when turning left or right (Fig. S5). To quantify turning, we extracted rotational ball movement (left and right) using its optic flow and convolved it with the fluorescence increase of GCaMP for each experiment and component. This convolution matched the temporal characteristics of the behavior to brain activity, and smoothed potential jitter from the ball. The speed of the rotational flow, which is proportional to the degree of turning, was on average 0.4 - 2 radian/s. The components that correlated with turning were reproducibly found across different flies based on similar position and morphology (Fig. 5A). These included the IPS-Y (inverse Y shape in the posterior slope) and LAL-PS (lateral accessory lobe and posterior slope) as previously mentioned in (Sophie Aimon et al., 2019) which could result from DNb02 and DNb01 for IPS-Y and LAL-PS, respectively (Table 2). These DNs were previously described by Namiki et al., who generated specific Gal4 lines for neuronal manipulation (Namiki et al., 2018; Robie et al., 2017). For instance, optogenetic activation of DNb01 induced twitching of the fly’s front legs consistent with a role in turning (Cande et al., 2018). For dopaminergic neurons, the components most correlated with turning could be matched with specific neurons such as neurons of the PPM2 cluster, i.e., PPM2-LW (PPM2-LAL-WED: PPM2-lateral accessory lobe-wedge (Mao and Davis, 2009))(Fig. 5A, right panels; Movie 5). By subtracting the coefficient of activity on the contralateral side from the coefficient on the ipsilateral side, we found that some of these turning components were activated differentially depending on whether the fly turned left or right (Fig. 5B).

**Figure 5:**
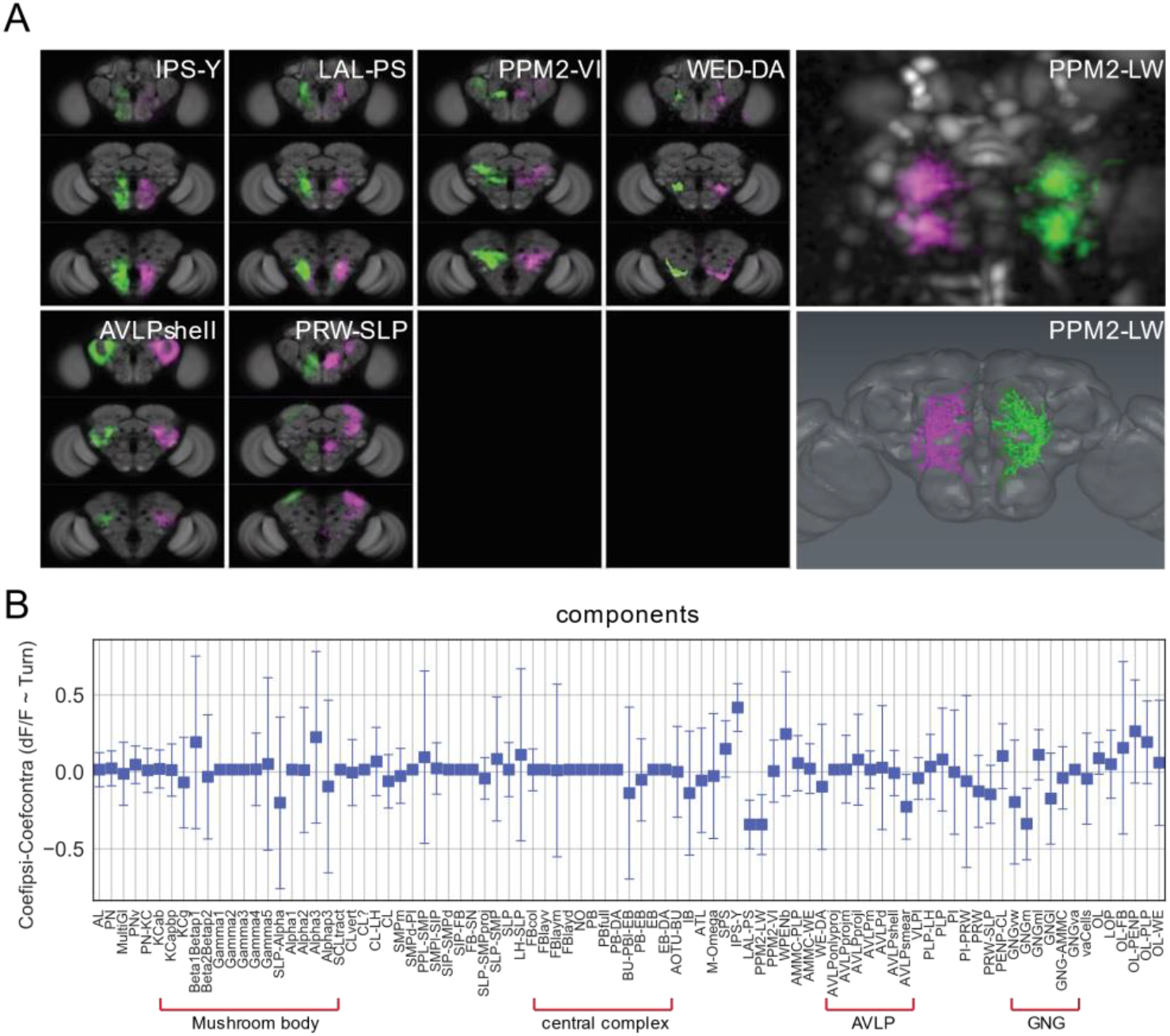
Turning activates specific components and neurons. (**A**) Examples of components present in both the left and right hemisphere labeled in different colors (magenta and green). Panels on the right present an example component that could be mapped to a single neuron. Upper right panel: Turning-correlated component, lower right panel: reconstruction of neuron that this functional component was mapped to. (**B**) Difference between the correlation coefficient (normalized ΔF/F) for turning on the ipsilateral side and the coefficient on the contralateral side is displayed as a function of the identified components. Positive and negative correlations correspond to components being active more during turn on the ipsi-lateral side than the contra-lateral side and the reverse, respectively. See Table 2 for definition of acronyms. N=58 flies of different genotypes, see table in methods for details.

These data show that part of the globally observed increase in neuronal activity during spontaneous walk corresponds to spontaneous turning behavior. As for forward walk, most components could be matched to candidate neurons (Table 2) for future functional analysis.

### Brain dynamics at transitions between rest and walk

So far, we have analyzed neuronal activity during walk, turn, flail, groom or resting. Next, we analyzed whole brain activity at the transition between rest and walk. Importantly, different GCaMP versions showed equally fast onset dynamics, non-distinguishable under our experimental conditions. To get a better temporal resolution, we only included datasets recorded at 30 Hz (with a maximum of +/- 30 ms error between behavior and brain activity) or faster for the entire brain in Fig. 6 (see all individual trials including data recorded at less than 30 Hz in Fig. S6). We normalized activity at the onset of walk and averaged trials. This analysis revealed a more complex picture of neuronal activity with some components being activated before the fly started to walk (Fig. 6).

**Figure 6:**
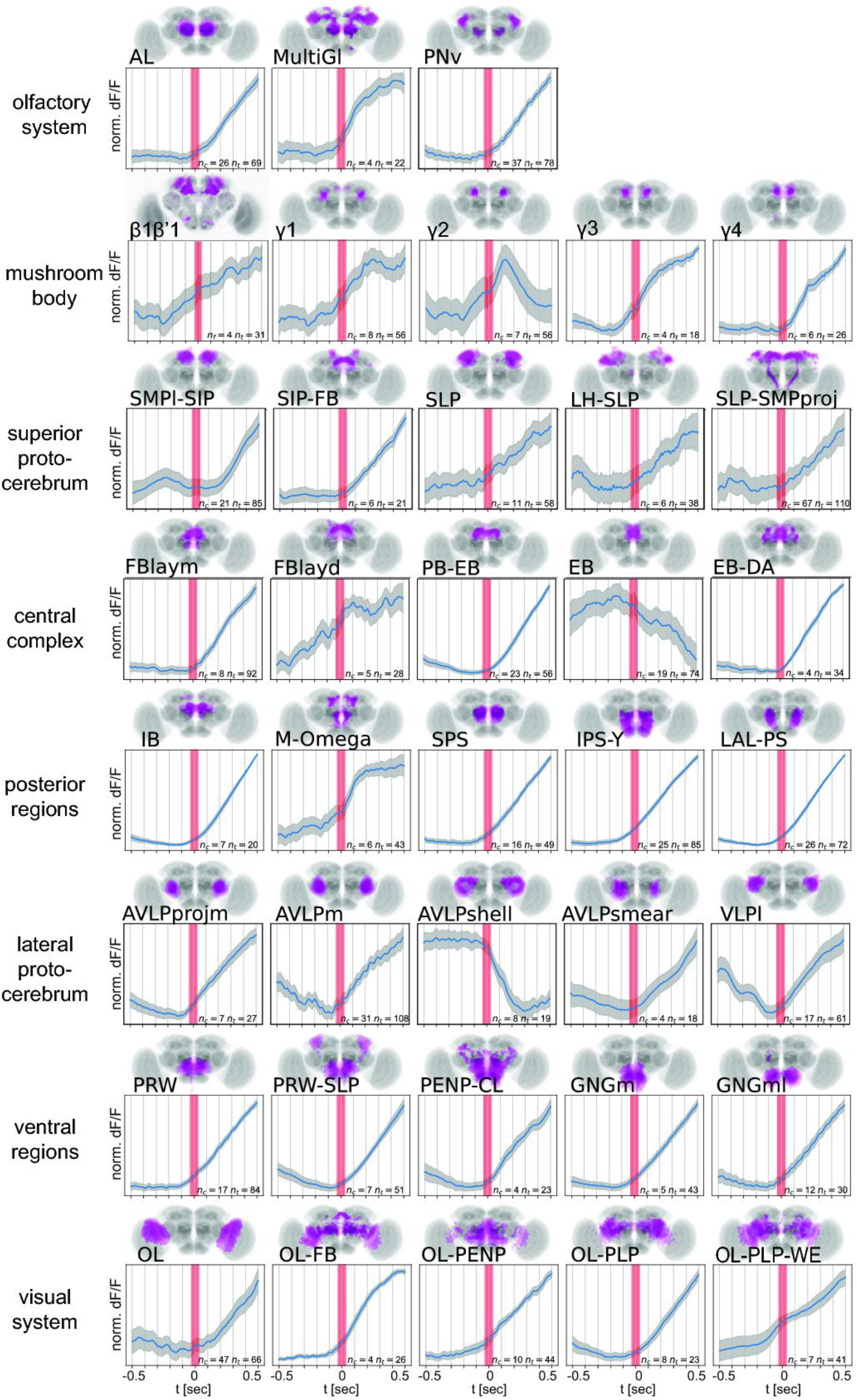
Walk induces activity in multiple functional components across the brain at start of walk. Walk-onset triggered average activity for pan-neuronal data in individual active components at 30 Hz or higher temporal resolution. Trials were normalized individually before averaging. Note that while most components are activated after walk onset, several components show activity before start of walk. See Table 2 for definition of acronyms. N=30 flies, see table in methods for details.

Components most reliably activated just before walk were regions around the esophagus such as the PRW, posterior slope (IPS-Y, SPS) and GNGm which activity started to increase ~100ms before walk onset. This is consistent with a role in movement initiation and the presence of many descending neurons in these regions (Lee et al., 2020). Note that activity before walk onset could also be due to preparatory movements that were not detected as walk (Ache et al., 2019). In addition, several compartments of the MB showed trials with some increase preceding walk (Fig. 6). Interestingly, while MB-compartment γ4 was active at start of walk, neighboring compartments such as γ3 could show an activity increase at ~200 ms before the animal started to move (Fig. 6, Fig. S6). Activity before walk onset was also visible in some trials in superior neuropile regions such as SMPm and SLP, which are directly connected to compartments of the MB. By contrast, the α-compartments showed little change in activity at walk onset (Fig. S6). These data are consistent with a role of specific MB-compartments in contributing to the control of motor output (i.e. γ1-3) as well as in integrating walk-related feedback signals (see also (Zolin et al., 2021)). Another brain area previously implicated in movement and navigation is the central complex (e.g., (Lyu et al., 2022; Seelig and Jayaraman, 2013)) comprising the protocerebral bridge (PB), fan-shaped body (FB), ellipsoid body (EB), and noduli (NO). Most components mapped to these brain areas were activated at walk onset but not before (Fig. 6, S6A) in line with the role of the central complex in navigation and orientation. In particular, the components PB-EB, connecting the PB and the EB, and EB-DA, comprising neurons in the EB and the lateral accessory lobe, became activated once the fly started to walk (Fig. 6, S6A). Of note, the component PB-EB corresponds to the previously described head direction cells shown to receive movement-related information during navigation (Lu et al., 2022; Lyu et al., 2022; Seelig and Jayaraman, 2015). EB-DA, found in data from dopaminergic neurons, shows among the strongest and most reliable activity during walk. These neurons were shown to be involved in ethanol-induced locomotion (Kong et al., 2010) and are also involved in sleep regulation (Liang et al., 2016). An exception from in the central complex were the ventral layers of the FB (FBlayv, Fig. S6) and the EB components from lines other than TH-gal4 that did not show an increase after walk onset (Fig. 6). The dorsal layer, FBlayd, showed some slight ramping up activity before walk-onset (Fig. 6) in line with its role in arousal and sleep homeostasis (Donlea et al., 2014). Distinct components mapped to the AVLP displayed very different activities related to walk. While anterior and medial components of AVLP increased in activity at walk onset, the AVLP component that resembled the shape of a shell (AVLPshell) displayed a very clear decrease in neuronal activity as soon as the fly started to walk (Fig. 6, S6A). A serotonergic neuron of unknown function called Trh-F-100082 by Chiang et al. might contribute to this activity pattern (Chiang et al., 2011). Several other components displayed more variable, and at times too variable dynamics between flies to detect a clear direction (Fig. S6).

As a complementary approach to the component analysis, and to further investigate putative differences in dynamics between neuron types, we next analyzed the timing of activity relative to walk onset of the different neuromodulatory neuron lines in anatomically defined brain regions (Fig. S7). Given the suggested importance of DANs for movement preparation and initiation in vertebrates (Coddington and Dudman, 2019), we next analyzed DAN activity more carefully (Fig. S7A). In most brain regions, DAN activity increased at the start of walk. In others such as in regions of the superior medial protocerebrum (SMP) and the mushroom body, the activity went up before the fly started to walk and stayed high during walk (Fig. S7A). Octopaminergic neurons were also mostly activated at walk onset (Fig. S7B). We observed complex activation patterns for serotonergic neurons (Fig. S7C). First, we observed a decrease in activity at walk onset for several brain regions including the AVLP, the PLP, MB calyx (MBCA) and the ICL. However, some regions showed an increase at walk onset (e.g., GNG, saddle (SAD) or SIP) presumably through distinct types of serotonergic neurons (Fig. S7C).

The observed activity patterns confirm and extend published neuronal manipulation data, where available, on sleep and locomotion (see above and discussion). Our data suggest that the activities of these different types of neurons with distinct functions in state or behavioral control contribute to the global change in brain activity elicited before and during walk. Our analysis provides an entry point to investigate how individual neurons or brain regions contribute to global brain states and which role this contribution to brain state plays in modulating behavior and sleep.

### Forced walk and forced turning recapitulates most activity

Our data suggest that walking induces a change of activity in most of the brain. But where does this activity come from? We hypothesized that in the extreme cases, activity could essentially originate from two opposite sites. First, the activity could arise initially in superior decision-making areas and then spread across the brain (top-down) or, second, activity is initiated by motor activity and proprioception (bottom-up) and then distributes to higher brain areas. In the latter case, the activity would originate in the VNC and move to the basal regions of the brain, i.e., the GNG, via ascending neurons (AN) (Chen et al., 2022; Tsubouchi et al., 2017). While our component analysis of data recorded at 30 Hz identified several putative ‘top-down’ components active before walk onset, most activity was more consistent with a ‘bottom-up’ scenario (Fig. 1, 6 and S6). In some trials, we observed that the activity detected during walk appeared to progress from areas at the base of the brain, i.e. GNG to more dorsal areas (Fig. S8D, see Movie 6 for an example trial).

We thus asked whether the walk-induced activity could be contributed by axon terminals of ANs. To this end, we expressed a synaptically-tethered GCaMP, syt-GCaMP6, under the control of a pan-neuronal driver and imaged whole brain activity during walk (Fig. 7A). Compared to cellular-GCaMP, syt-GCaMP activity was very strong in the GNG, AMMC and AVLP, the regions receiving input from the VNC (Fig. 7A) (Tsubouchi et al., 2017).

**Figure 7:**
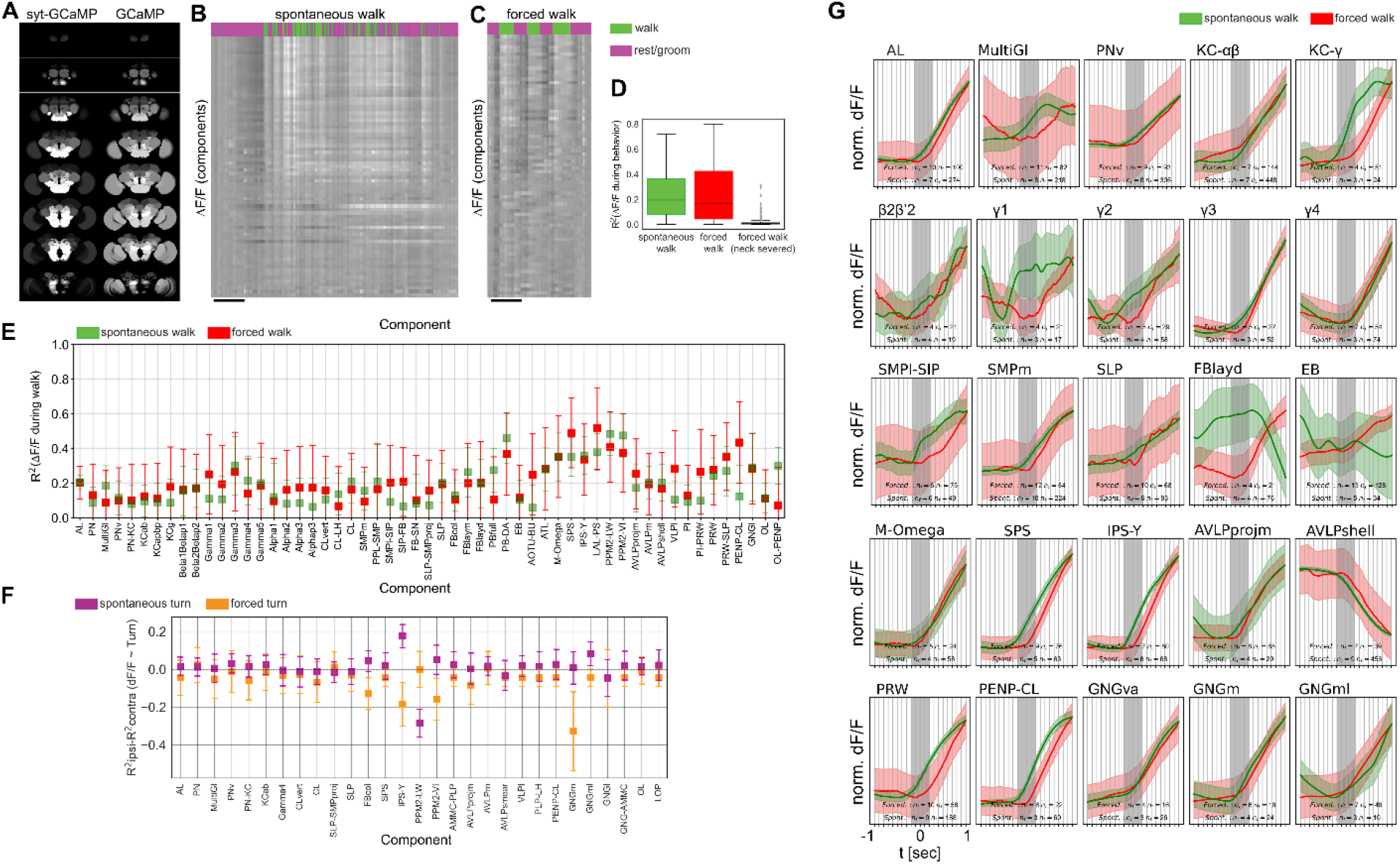
Forced and spontaneous walk elicit highly similar global brain activity. (**A**) Z-stacks of map of brain regions activated by walk in flies expressing cytosolic GCaMP (pan-Gal4;UAS-GCaMP6m) or synaptically tagged GCaMP (pan-Gal4;UAS-GCaMP6m) (**B**) Time series of components during spontaneous walk or rest (magenta vs. green). (**C**) Time series of active components during forced walk or forced rest (magenta vs. green). (**D**) R^2^ for behavior regression at different conditions (all regions were pooled, spontaneous, forced, forced with severed connection between VNC and brain) (**E**) R^2^ for behavior regression for different active components for forced vs. spontaneous walk. (**F**) R^2^ difference between brain region activity during forced turn on the ipsilateral side and forced turn on the contralateral side (orange). Only data with n>=4 components were analyzed. Lilac shows the difference for spontaneous turns. N=58 flies for spontaneous walk/turn of different genotypes and N=26 flies for forced walk/turn, see table in methods for details. Box plots show: center line, median; box limits, upper and lower quartiles; whiskers, 1.5x interquartile range; points, outliers. (**G**) Comparison between activity at walk onset for spontaneous (green) and forced (red) walk, for additional components. Individual trials were normalized and averaged for each component. Shaded regions represent the trail SEM, nf: number of flies, nt: total number of trials.

Since such axonal activity, or a fraction of it, could potentially also originate from top-down projections, we carried out another experiment and compared spontaneous, self-induced walk to forced walk. We argued that if the broad activation indeed comes from proprioception of walking or related sensory input from the legs, we should also observe a global activation when the fly is forced to walk rather than decides to walk spontaneously. To test this idea, we placed a walking substrate controlled by a motor under the fly legs (see Movie 7 and methods). We forced the flies to walk by turning this motor on and off at a speed range between 1.5 to 6 mm/s. Spontaneous walk in these flies elicited the typical whole brain activity as described above (Fig. 7B,D). Remarkably, forced walk also induced a highly similar global activity across the brain (Fig. 7C,D, Movie 7), and across components (Fig. 7E). In addition, flies with a surgically severed connection between brain and VNC walked on the treadmill while showing hardly any activity in the brain (Fig. 7D, Movie 8) consistent with the hypothesis that the brain is not necessary for forced walk and walking signals driving whole brain state change originate in the VNC. Importantly, these observations were not only true for pan-neuronal data but also for dopaminergic, octopaminergic and serotonergic neuronal subsets (Fig. S8A-C). While we did observe some variation between forced and spontaneous walk in the R^2^-value for walk, no significant difference was found for any of the brain regions (Fig. S8A-C) further suggesting that walking, whether spontaneous or forced, elicits a highly similar global state change of the brain.

By contrast, differences between left and right hemispheres in flies that were forced to turn by 0.3 to 2 radian/s significantly differed from lateral differences in flies that turned spontaneously (Fig. 7F). We subtracted the R^2^-value for turn of the contralateral side of the brain from the corresponding value of the ipsilateral side and compared these values for individual components. We found that some components switched sign (i.e., IPS-Y) while other components that showed little lateralization during spontaneous turns showed strong differences for forced turns (i.e., PPM2-VI, GNGm, FBcol) (Fig. 7F).

Given our observation that certain brain regions appear to be activated shortly before walk onset (see Fig. 6), we next compared the activity around walk onset in spontaneous and forced walk more carefully (Fig. 7G, S8E). Most active components appeared very similar as expected based on the analysis above (Fig. 7E). In fact, the normalized activity traces in several components overlapped between forced and spontaneous walk (e.g. γ4, m-omega, Fig. 7G, S8E). This included several components located in the GNG consistent with our hypothesis that the activity observed in GNG originated primarily from the VNC and walking movements themselves. Importantly, the components with activity prior to spontaneous walk were activated later in forced walk, namely only when walk had been triggered (i.e. components in the γ lobe of the mushroom body, posterior slope, prow, and dorsal layer of the fan-shaped body Fig. 7G, S8E) consistent with a motor command function of these brain regions.

Taken together, these data are consistent with a model wherein a substantial part of the whole brain activity observed during walk is induced by walk signals from the VNC and/or by walk-related proprioceptive stimulation and not by top-down activity of higher brain centers. The finding that the same components are active when the fly decides to walk spontaneously and when it is forced to walk suggests that components with motor command contribution also receive strong feedback from walk. As the overall brain activity increase was not larger during forced walk as compared to spontaneous walk, our data also suggests that brain activity does not represent a mismatch, or error signal, between actual and predicted proprioceptive feedback during walk. However, more detailed analysis of the lateralization of activity during spontaneous *vs.* forced turns revealed some local differences, which could be consistent with error signals. Finally, our analysis revealed several brain regions such as the posterior slope and, in some trials, the γ-compartments that are active prior to walk onset in spontaneously walking flies but not during forced walk suggesting that these brain areas direct or prepare walk and receive feedback when the animal has started to walk.

## Discussion

Work over the last years has revealed that locomotion and movement influence the activity of neurons in many brain areas and organisms (Busse et al., 2017; Kaplan and Zimmer, 2020). Importantly, motor activity modulates not only local activity in specific brain regions, instead it appears to change the global state of the brain. Where this activity originates from, how it spreads, and what it means for the animal is still debated (Kaplan and Zimmer, 2020). Using a tethered fly preparation and fast *in vivo* whole brain imaging, we showed that movement elicits a global change in brain activity during spontaneous as well as forced walk. Our data further suggested that walk activates several different classes of neurons including excitatory (cholinergic) neurons, inhibitory neurons as well as aminergic, modulatory neuron types. Except for serotonergic neurons which are inhibited during walk in some brain areas, we observed neuronal activation across all brain regions at the start of and during walking, but not during grooming or resting. Using PCA/ICA, we mapped neuronal activities to discrete functional components, which we assigned to specific smaller subregions and in some cases even to single neurons by aligning the activity data to anatomical images. For instance, we pinpointed specific neurons activated during turning. Based on our data and analysis, we propose that locomotion activates the brain by sending movement and proprioceptive information to the base of the brain (the GNG) from where it activates all brain regions. These data and analysis inspire testable hypotheses and provide a resource of potential neural substrates involved in the perception or control of walking.

### Role and origin of broad activation during ongoing behavior

One important concept to explain the role of behavioral state-dependent neural modulation is referred to as ‘Active sensation’ (Busse et al., 2017). Essentially, ongoing movement can shape how neurons respond to visual, somatosensory and other sensory stimuli (Chapman et al., 2018; Cruz et al., 2021; Fenk et al., 2021; Henschke et al., 2021; Wolpert et al., 2011). For example, extracellular recordings from V1 neurons of mice walking on a ball showed that evoked visual responses differed during movement as compared to neurons of still animals (Dadarlat and Stryker, 2017). Similarly, fly visual neurons respond stronger stimuli during active walk or flight (Chiappe et al., 2010). Our data indicate that whole brain activity increase is elicited with walk onset and maintained afterwards (see Figure 6). Our analysis furthermore revealed brain areas and select neurons that respond to turning (see Fig. 5). These observations support the conclusion that movement specific, proprioceptive and other sensory information reach the brain and modulate brain activity widely. Such information could serve multiple purposes from uncoupling of sensory-to-motor information, i.e. own movement *vs.* movement of environment, to learning of complex movements. Brainwide neural and glial responses appear to enable behavioral flexibility in zebrafish (Mu et al., 2020). By coupling_sensory processing and behavior in a closed loop, brain activation during ongoing behavior could resemble a form of working memory, in part for learning to improve future behavior or to relate body-movement to environmental information (Lu et al., 2022).

Our combined data suggests that while many regions are active during walk, they play different roles. First, activity before walk onset that was not present in forced walk likely pinpoints neurons involved in triggering walk. Second, these regions are also active during forced walk, albeit with a temporal delay, indicating that they receive feedback that walk has indeed been triggered. Third, temporal and spatial dynamics are contributed by neurons producing all major classes of neurotransmitters or neuromodulators. This combined activity is consistent with several functions of the observed global brain state change: motor command, motor maintenance and error signals, arousal, as well as motor learning.

Our results are most consistent with a model where proprioceptive, walking and leg sensory information are sent from the VNC into the GNG at the base of the fly’s brain. So, how is walking related information relayed to the brain? Tuthill and colleagues recently identified some of the presumably many neural substrates in the VNC that receive, process and relay proprioceptive sensory information from the legs to the CNS (Agrawal et al., 2020; Mamiya et al., 2018). Their findings provide strong support for an important role of proprioception in movement and locomotion control in the adult fly. This information is transmitted by ascending neurons from the VNC to the central brain. So far, relatively little is known regarding the type, connectivity and function of the likely dozens or more ascending neurons in *Drosophila,* but screening approaches and advanced imaging techniques have shed light on some of them (Allen et al., 2020; Chen et al., 2018; Sen et al., 2019).

While grooming or forced walk on a treadmill do not require an active brain, lesions of the neck connectives as we have carried out dramatically decrease spontaneous walking in locusts (Kien, 1990a, 1990b) indicating that at least initiation of walk can be dependent on the brain. On the other hand, a cat with a severed spinal cord, like our fly with a severed VNC, maintains a highly coordinated walk pattern when forced to walk on a treadmill (Afelt, 1974). Thus, coordinated walk appears to mainly depend on central pattern generators in the spinal cord or VNC and is largely independent of brain input. Surprisingly, whole brain activity induced by spontaneous walking was similar to the activity we observed by forcing the animal to walk on a rotating rod (see Fig. 7). This activity disappeared when the connection between VNC and brain was disrupted. It is conceivable that a similar effect would be observed in a cat or a mouse being forced to walk on a treadmill with a severed spinal cord. This result and our finding that the activity first induced by walk in the GNG stems from axons are consistent with the interpretation that walk itself and not top-down motor control is responsible for the majority of activity observed in actively moving animals’ brains.

Importantly, however, we identified several brain components and small subregions, for instance in some trials in the MB, that were activated hundreds of milliseconds before the fly started to walk (see Fig. 7). This activity was delayed in forced walk and started only when the fly had started to move. This finding suggest that MB γ-compartments 1-3 are involved in triggering walk and receive behavioral state feedback while compartments γ4 and γ5 are only activated when the fly has already decided to move. Similar interpretations might apply to regions of the central complex (see Fig. 7). Considering these findings in the context of what we already know about the MB, our data suggests that compartments implicated in aversion (i.e. γ1 and 2) are capable or involved in triggering walk behavior, while compartments involved in appetitive behaviors merely receive the information that the animal walks. More generally, our data indicate that regions triggering walk are also highly sensitive to walk itself suggesting a closed-loop between walk initiation and walk maintenance, possibly including factors such as speed, detailed movement etc. Such an organization ensures the close coordination between what the animals plans and what it already does.

Our data provides an entry point to relate anatomical connectivity to activity in widespread neural networks (Aimon and Grunwald Kadow, 2019). Whole brain connectomics in several small organisms including *Drosophila* has shown that neural networks extend across the entire brain with many pathways not predicted by single neuron or small motif imaging or manipulation of individual neurons (Li et al., 2020; Scheffer et al., 2020; Zheng et al., 2018). Wholebrain imaging, as previously demonstrated in other model animals such as larval zebrafish, is a powerful method for observing these brainwide networks to study their contribution to behavior and ultimately to local activities within specific neurons or brain regions. Whole brain data can be used to build functional connectomes allowing speculations about how information underpinning behavior travels throughout a whole brain. While such data can be generated for other animals including humans, the fly (along with C. elegans and in the near future zebrafish) currently provides the important advantage of being able to combine such activity maps with highly detailed anatomical maps from light microscopy with cellular resolution and whole brain EM connectomics with synaptic resolution. Ultimately, such data could be used to generate precise models of how recorded neural activity spreads through a brain. For instance, DNs receive input from regions of the brain that are innervated by outputs from higher brain regions such as the MB and central complex (CC) (Hsu and Bhandawat, 2016). DN and AN innervation of the GNG is consistent with an important role of the GNG in motor control and motor feedback integration (Chen et al., 2022; Tsubouchi et al., 2017). How anatomical connectivity relates to activity in widespread neural networks is not clear.

### Walk elicits differential activities in neuromodulatory neurons

Perhaps not surprisingly, neuromodulatory systems participate and show signatures of ongoing behavior in the adult fly brain (see Fig. 3). Dopaminergic and octopaminergic neurons are broadly activated when the fly walks compared to rest or groom (see Fig. 3 (Aimon et al., 2019; Siju et al., 2020)), while serotonergic neurons show more complex activation patterns and timing with areas such as the AVLP being inhibited during walk (see Fig. 3 and S6).

This data suggest that relationships between the activity of neuromodulatory neurons and locomotion are broadly conserved across species. Dopamine neurons are activated during locomotion in mammals, with some neurons reporting ongoing behavior (Howe and Dombeck, 2016)while others have activity increase preceding behavior onset (Coddington and Dudman, 2019). We find that although the majority of fly dopamine neurons are activated during ongoing behavior, some dopaminergic neurons are activated hundreds of milliseconds before the onset of walk in some trials (e.g. γ1-3 compartments of the mushroom body, see Fig. 6). Norepinephrine/octopamine is also a key neuromodulatory systems involved in arousal in a variety of species (Berridge, 2008), with reports of an increase in activity of these neurons during locomotion in mammals (Gray et al., 2021; Kaufman et al., 2020; Xiang et al., 2019), also consistent with our observations in flies. Interestingly, serotonergic neurons show mixed roles in the control of motor behavior in different vertebrate species (Dayan and Huys, 2015; Flaive et al., 2020; Vitrac and Benoit-Marand, 2017), as well as in our fly data. In particular, serotonin has been implicated in behavioral inhibition (e.g. patience (Doya et al., 2021)) and as an opposing system to dopamine (Dayan, 2012). This is consistent with our observation that serotonergic neurons in the AVLP are inhibited during walk. Similarly, serotonergic neurons in the fly VNC are also inhibited during walking behavior as previously reported (Howard et al., 2019). However, we also found some serotoninergic neurons activated during walk, suggesting that, as for mammals, serotonin neurons are strongly heterogeneous(Dayan and Huys, 2015). Our study thus suggests that the activity of neuromodulatory neurons during locomotion is broadly conserved between mammals and insects despite their evolutionary distance.

These neuromodulators could be involved in the closed loops linking sensory processing and behavior. In vertebrates, basal ganglia and brainstem aminergic neurons affect the cortico-basal ganglia-thalamic loops. A disruption of these loops can result in a loss of motor control (Vicente et al., 2020). Such loops likely exist in insects, too. For example, several octopaminergic neurons connect lower brain regions to the MB (Busch et al., 2009). Dopaminergic neurons innervate central complex and MB (Mao and Davis, 2009). In line with this, we observe a strong activation of dopaminergic and octopaminergic neurons during walk. Serotonin has varied roles in vertebrates and regulate various types of motor behaviors including feeding, aggression and larval locomotion in insects (e.g., (Aonuma, 2020; Helfrich-Förster, 2018; Ngai et al., 2019; Schoofs et al., 2018, 2014)). The AVLP, we identify as a key region where serotonin neurons are inhibited during walk (see Fig. 3 and 6) receives input from ascending neurons from the VNC conveying somatosensory information from the legs (Tsubouchi et al., 2017). Interestingly, calcium imaging revealed a spatial map for the AVLP and WED with neurons responding primarily to movement of fore-, mid- or hindlegs (Chen et al., 2022; Tsubouchi et al., 2017). The walk-related role of the strongly inhibited region we termed AVLPshell has not been studied yet, to our knowledge. The functional component maps matched to anatomical templates should thus be helpful in identifying neurons within these regions and their respective functions during walk.

### Advantages and limitation of the method

Light field microscopy (Levoy et al., 2006) has several advantages and limitations, which complement other existing imaging methods. First, the light field speed –significantly higher temporal resolution as compared to sequential scanning methods– allowed us to record whole brain activity at the same time as fast behavior such as walk. Even though GCaMP dynamics and limitations in signal-to-noise ratio rarely permitted us to resolve single action potentials, it helped us detect differences in temporal dynamics and pinpoint brain areas involved in triggering spontaneous walk as opposed to merely responding to it. Second, the spatial resolution of lightfield imaging is inferior to that of confocal or volumetric multiphoton imaging. Although these methods also do not allow resolving single neurites with pan-neuronally expressed sensors, lightfield could in principle make it even more difficult to detect their activity. On the other hand, capturing the whole volume simultaneously makes it easier to unmix signals as we showed with our PCA/ICA approach, which partially compensates for the lower spatial resolution. Third, our data is based on observations without genetic or functional manipulation of neurons or circuits. Excitation and inhibition of single neurons are being carried out very frequently in *Drosophila* thanks to its unique genetic tools. We believe that our data complements previous and future functional studies as imaging or manipulation of individual neurons provides only limited insights into the role and effect of a neuron in the complex dynamic neural networks in which they are embedded. In the future, a combination of single neuron manipulation and wholebrain imaging will likely lead to unexpected insights into the relationship of a neuron and a specific behavior. Fourth, we have observed subtler differences between flies that are obvious from individual experiments but difficult to capture quantitatively across a population of animals. These differences might be resolvable by greatly increasing the number of experiments. Given the technical difficulty of the preparation method, reaching high animal numbers will be extremely challenging but perhaps possible in the future. Finally, together with the now available whole brain EM connectome, our data provides a timely resource for the community of fly neuroscientists interested in linking neuronal activity to behavior.

### Conclusions

We provide an overview of brain activity during simple behaviors in *Drosophila*. As for other animals, *Drosophila* brain activity is globally correlated with locomotion leading to global change in brain state. However, our results challenge the assumption that most of the activity is related to decision-making, top-down motor control, or prediction error detection from sensory feedback and instead suggest that walk itself and somatosensory bottom-up stimuli are largely responsible. By using a combination of pan-neuronal and specific neuron line imaging, we shed light on the nature of neurons and their location in the brain that respond so strongly to behavior. Altogether, our data provide a novel resource for further generating new hypotheses regarding the brain-behavior-loop and for dissecting the neural circuits underpinning it.

## Materials and Methods

### MATERIALS

| Experimental models: Organisms/strains |  |  |
| --- | --- | --- |
| UAS-GCaMP6s | Bloomington<br>Drosophila Stock<br>Center | BDSC_42749 |
| UAS-GCaMP6m | Bloomington<br>Drosophila Stock<br>Center | BDSC_42750 |
| UAS-GCaMP7s | Bloomington<br>Drosophila Stock<br>Center | BDSC_79032 |
| UAS-GCaMP6f | Bloomington<br>Drosophila Stock<br>Center | BDSC_42747 |
| UAS-GCaMP7f | Bloomington<br>Drosophila Stock<br>Center | BDSC_79031 |
| UAS-syt-GCaMP6s | Vanessa Ruta | N/A |
| GMR58E02-Gal4 | Bloomington<br>Drosophila Stock<br>Center | BDSC_41347 |
| Th-Gal4 | Bloomington<br>Drosophila Stock<br>Center | BDSC_8848 |
| Tdc2-Gal4 | Bloomington<br>Drosophila Stock<br>Center | BDSC_9313 |
| Trh-Gal4 | Bloomington Drosophila Stock Center | BDSC_38388 |
| Ddc-Gal4 | Bloomington Drosophila Stock Center | BDSC_7009 |
| TH, 58E02-Gal4 | Siju et al. | N/A |
| TH, DDC-Gal4 | This paper | N/A |
| Vglut-Gal4 | Bloomington Drosophila Stock Center | BDSC_24635 |
| Cha-Gal4 | Bloomington Drosophila Stock Center | BDSC_6798 |
| Nsyb-Gal4 | Bloomington Drosophila Stock Center | BDSC_51635 |
| GMR57C10-Gal4 | Bloomington Drosophila Stock Center | BDSC_39171 |
| Gad1-Gal4 | Bloomington Drosophila Stock Center | BDSC_51630 |
| **Software and algorithms** |  |  |
| Python 3 | Python Software Foundation | https://www.python.org |
| ImageJ | Schneider et al., 2012 | https://imagej.nih.gov/ij/ |
| Fiji | MPI-CBG | https://fiji.sc/ |
| MATLAB | MATLAB | [mathworks.com](https://www.mathworks.com) |
| Analysis code | GitHub | https://github.com/sophie63/Aimon2022 |

#### Fly preparation for imaging

We used one to four days old female flies raised at 25 °C. Most flies were starved 24 or 48h with a water only environment, and we clipped their wings at least one day in advance. Experiments were performed in the evening peak of circadian activity (ZT0 or ZT11) and we heated the room to ~28 °C during the experiment. In total, we recorded brain activity and behavior from 84 adult female flies.

We prepared the flies as described in detail in (Woller et al., 2021). Briefly, we fixed a fly to a custom-designed 3D printed holder, so as to allow access to the whole posterior side of the head while the legs were free to move. We added saline (103 mM NaCl, 3 mM KCl, 5 mM TES, 8 mM trehalose 2 H_2_O, 10 mM glucose, 26 mM NaHCO_3_, 1 mM NaH_2_PO_4_, 2.5 mM CaCl_2_·2 H_2_O, 4 mM MgCl_2_·6 H_2_O) and dissected the cuticle, muscles, and air sacks at the back of the head to expose the brain.

#### Walk substrates

For studying spontaneous walk, we used two types of small balls. One was an air-supported ball as previously described (Sayin et al., 2019). As we wanted to make sure the walk was initiated by the fly rather than erratic movement of the ball, we also used small styrofoam balls that were held by the fly.

As treadmill for studying forced walk and turn, we used small motors (DC 6V gear motor with long M3 x 55mm lead screw thread output shaft speed reducer Walfront Store, www.amazon.de), covered with self-curing rubber (from Sugru) to provide a smoother surface. The speed for forced walk was between 1.5 and 6 mm/s. The rotation speed for forced turn was 0.3 to 2 rad/s.

#### In vivo light field imaging

Fast volumetric imaging was performed using light field imaging – in which a microlens array separates rays from different angles to give information on depth – and was carried out as previously described in detail by (Aimon et al., 2019). A few datasets were previously published in Aimon et al. (Aimon et al., 2019) and source data (http://dx.doi.org/10.6080/K01J97ZN), with a microscope equipped with a 20x NA1.0 objective. Most data were obtained with a light field microscope constituted of a Thorlabs Cerna system with a Leica HC FLUOTAR L 25x/0.95 objective and an MLA-S125-f12 microlens array (Viavi). The microlens array was placed on the image plane, while the camera imaged the microlens array through 50 mm f/1.4 NIKKOR-S Nikon relay lenses. The light field images were recorded with a scientific CMOS camera (Hamamatsu ORCA-Flash 4.0). The volumes were reconstructed offline, using a python program developed by (Broxton et al., 2013)(*76*) and available on github: https://github.com/sophie63/FlyLFM.

Given that the maximum recording speed depended on the expression of the Gal4-line, the UAS-reporter, and each individual fly preparation, i.e. the limiting factor was the signal-to-noise ratio for lines with low expression, we started recording at 2 or 5 Hz for the first experiment. If the quality of recording suggested that higher speeds were possible, we increased recording speed to a maximum of 98 Hz. If the viability of the fly allowed for it, we recorded experiments on air-supported and Styrofoam balls in the same fly.

| Gal4 | UAS<br>-<br>GCa<br>MP | # of<br>flies,<br>spontan<br>eous<br>Walk<br>(flail,<br>groom) | flailing | grooming | frame rate in Hz<br>(# of experiments) | Walking<br>Substrate<br>(# of<br>experiments) | # flies<br>of flies<br>forced<br>walk |
| --- | --- | --- | --- | --- | --- | --- | --- |
| total | total | 58 | 9 | 6 | 98(8), 50(21), 20(11),<br>10(9), 5(30), 2(2) | Air-supported ball<br>(33), styrofoam<br>ball (54) | 26 |
| Nsyb | 7f | 1 | 1 | 1 | 50(1) | Air-supported ball |  |
|  | 7s | 1 | 1 |  | 5(1) | styrofoam ball | 1 |
|  | 6m | 6 | 5 | 2 | 50(8), 20(5), 10(3), 5(4) | Air-supported ball (16), styrofoam ball (5) | 3 |
|  | 6f | 3 |  | 1 | 98(2), 50(1) | Air-supported ball (2), styrofoam ball (1) |  |
|  | 6s | 1 | 1 |  | 50(1) | styrofoam ball(1) | 2 |
|  | total | 12 | 8 | 4 | 98(2), 50(11), 20(5), 10(3), 5(5) | Air-supported ball (19), styrofoam ball (8) | 6 |
| R57C10 | 7s | 1 |  | 1 | 98(2) | styrofoam ball(2) |  |
|  | 6s | 3 | 1 | 1 | 50(1), 5(1), 20(1) | styrofoam ball(3) | 3 |
|  | total | 4 | 1 | 2 | 98(2), 50(1), 20(1), 5(1) | Air-supported ball (0), styrofoam ball (5) | 3 |
| Cha | 6m | 4 |  |  | 5(4) | styrofoam ball (4) |  |
|  | 6f | 1 |  |  | 98(1) | Air-supported ball (1) |  |
|  | total | 5 |  |  | 98(1), 5(4) | Air-supported ball (1), styrofoam ball (4) | 0 |
| Vglut | 6m | 5 |  |  | 20(2), 10(3), 5(4) | Air-supported ball (3), styrofoam ball (6) |  |
|  | total | 5 |  |  | 20(2), 10(3), 5(4) | Air-supported ball (3), styrofoam ball (6) | 0 |
| Gad | 6m | 5 |  |  | 98(2), 20(1), 5(5), 2(1) | styrofoam ball (9) |  |
|  | total | 5 |  |  | 98(2), 20(1), 5(5), 2(1) | Air-supported ball (0), styrofoam ball (9) | 0 |
| TH/DDC | 6m | 5 |  |  | 50(2), 5(1), 10(1) | Air supported ball (3), styrofoam ball (2) | 4 |
|  | 6f | 6 |  |  | 50(5), 98(1) | Air supported ball (2), styrofoam ball (3) | 1 |
|  | total | 11 |  |  | 98(1), 50(7), 10(1), 5(1) | Air-supported ball (5), styrofoam ball (5) | 5 |
| Trh | 6m | 3 |  |  | 50(3), 5(3) | Air supported ball (1), styrofoam ball (6) |  |
|  | 6f | 3 |  |  | 10(2) | Air supported ball (2), styrofoam ball (1) |  |
|  | 6s | 3 |  |  | 5(3) | styrofoam ball(4) | 6 |
|  | total | 9 |  |  | 50(3), 10(2), 5(6) | Air-supported ball (3), styrofoam ball (11) | 6 |
| Tdc2 | 6s | 7 |  |  | 20(2), 5(4), 2(1) | Air supported ball (2), styrofoam ball (6) | 6 |
|  | total | 7 |  |  | 20(2), 5(4), 2(1) | Air-supported ball (2), styrofoam ball (6) | 6 |

#### Behavior recording and scoring

We imaged the fly and substrate movements using infra-red illumination and two small cameras FFMV-03M2M from Point Grey triggered by the fluorescence recording camera to ensure time alignment between fluorescence and behavior.

Walking, flailing, and grooming were obtained by measuring the optic flow from the Movies of the ball or by analyzing the movement of the fly’s legs using the “optic flow” plugin in FIJI. For turning or rotational speed (rad/s), the sum of left or right optic flows was not binarized. The behavioral time series were then convolved with the single spike response of the GCaMP version used for the experiment and subjected to the same deltaF/F procedure as the fluorescence time series (see below).

#### Pre-processing

Reconstructed volumetric fluorescence data was pre-processed by first correcting for movement using 3Dvolreg from AFNI (https://github.com/afni/afni). In Matlab, we then calculated the deltaF/F for each voxel by subtracting and dividing by the signal averaged for 4000 time points. We finally decreased noise with a Kalman filter (from https://www.mathworks.com/matlabcentral/fileexchange/26334-kalman-filter-for-noisy-movies) with a gain of 0.5.

We generated summary movies by maximum projecting the ΔF/F volumes and combining these to the behavior.

#### Alignment to template

We aligned the functional data to the anatomical template JRC2018 (https://www.janelia.org/open-science/jrc-2018-brain-templates) using the landmarks registration plugin with ImageJ (as described in http://imagej.net/Name_Landmarks_and_Register), with the landmarks found in https://github.com/sophie63/Aimon2022/blob/main/Registration/SmallJRC2018Template.points. allowed finding the anatomically defined regions covered by the functional regions (see Table 2) and find candidate neurons using Flycircuit (http://www.flycircuit.tw/) or Virtual Fly Brain https://v2.virtualflybrain.org data bases.

### STATISTICAL ANALYSIS

Statistics were performed in python with code freely available on https://github.com/sophie63/Aimon2022. To compare fluorescence time series (normalized by the absolute maximum value per fly) and behavioral time series, we used a simple regression model: TSfluo ~ BehaviorRegressor, solved with the ordinary least square fit function of the python statsmodels package. For each time series (either regional averaged intensity or the PCA/ICA component), this provided a fraction of variance explained by the behavior (R^2^), and the sign and strength of the correlation (coefficient). We compared these values with pair-wise tests using two-sided Mann-Whitney non-parametric tests with a Bonferroni multiple comparison correction. We used a more complex linear model to evaluate the effect of variables of interest (behavior, brain region, neural type) while explaining away confounds (GCaMP version, exact pan neuronal Gal4): R2 ~ Behavior+RegionNames+Gal4+UAS and Coef ~ Behavior+RegionNames+Gal4+UAS. We then plotted the coefficients + intercept, and 95% interval of coefficient + 95% interval of the intercept to compare the effect of the variables to zero. Flies were recorded over prolonged periods of times at different imaging speeds and during different behaviors. See table above for details on number of individual flies and types of experiments per fly.

#### PCA/ICA

To obtain functional maps of the fly brain, we performed PCA and ICA as described previously (Sophie Aimon et al., 2019; Beckmann and Smith, 2004)(*78*). Briefly, SVD (singular value decomposition) was used a first time to find the level of noise and normalize voxels by their noise variance. SVD was performed a second time on this normalized data resulting in maps and time series for principal components. The principal component maps were unmixed using ICA to obtain localized regions. The same matrix was used to unmix the time series.

## Acknowledgments

We are very grateful to Marta Costa and Kei Ito for sharing data, images, and knowledge during the course of this study. We also thank Francisco Rodriguez-Jimenez, Paul Bandow, Subhadarshini Parhi and Kunhi Purayil Siju for help with data analysis.

## Author contributions

Conceptualization: SA, IGK

Methodology: SA, IGK

Software: SA

Validation: SA, KC

Formal analysis: SA

Resources: IGK, SA

Data curation: SA, KC

Project administration: IGK, SA, JG

Funding acquisition: IGK, SA, JG

Visualization: IGK, SA

Supervision: IGK, SA

Writing—original draft: IGK, SA

Writing—review & editing: IGK, SA, JG, KC

## Data and materials availability

All original code is publicly available on Github. Original data and any additional information required to reanalyze the data reported in this paper are available upon request.

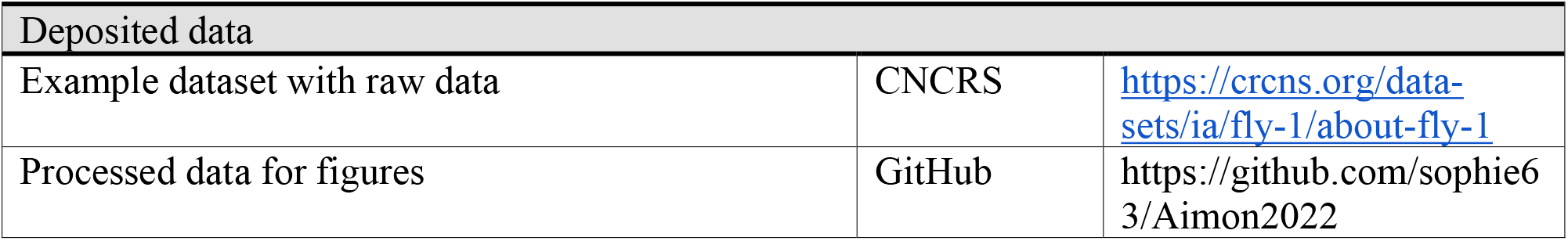

