## Supplementary figures and legends for "Combined patterns of activity of major neuronal classes underpin a global change in brain state during spontaneous and forced walk in *Drosophila*"

### Aimon et al., supplementary figures

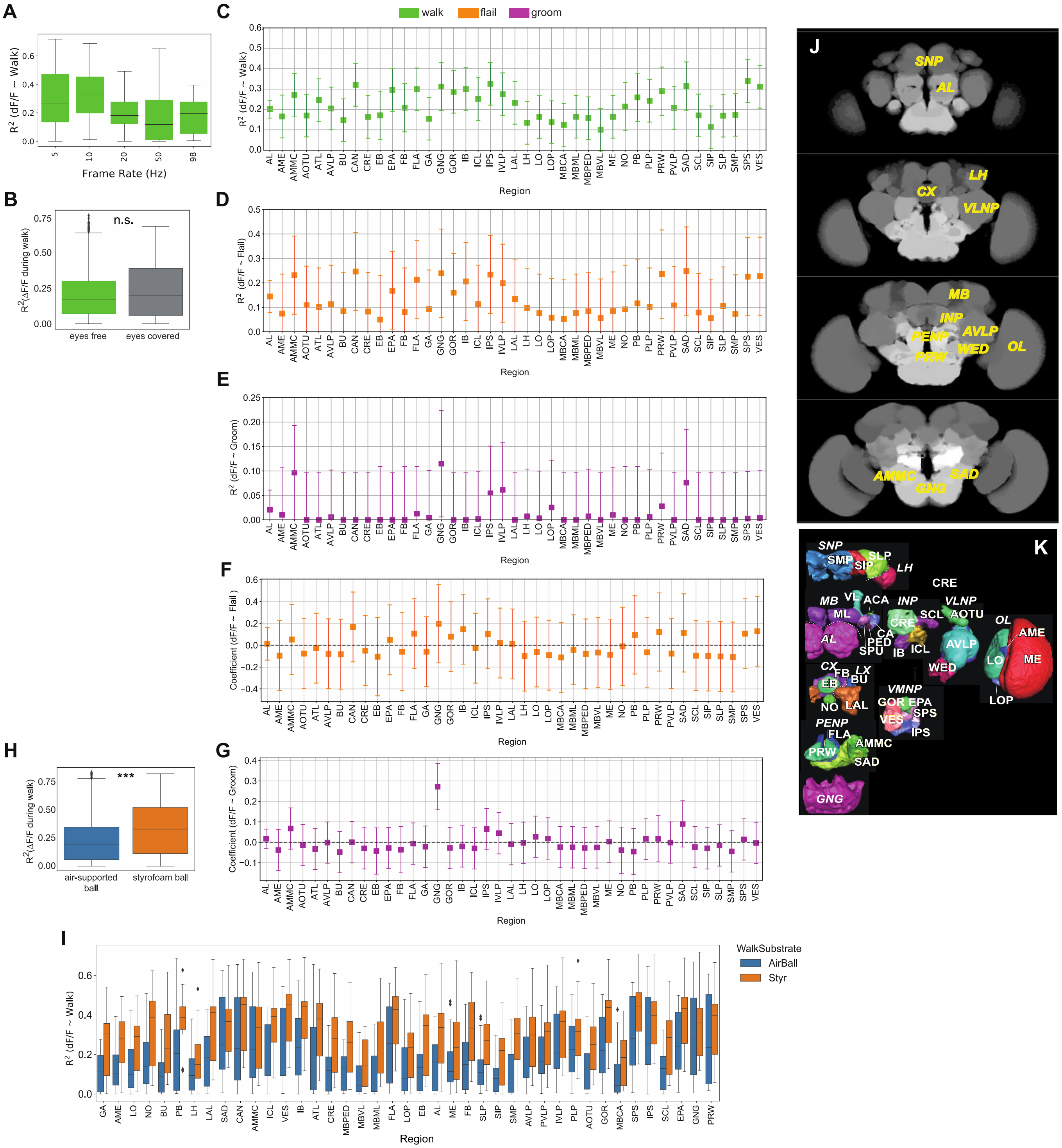

#### Fig. S1

(A) $R^2$ during walk at different recording frequencies for pan-neuronally expressed GCaMP6f (all regions were pooled). No significant difference was found for walk between the different frequencies. Box plots show: center line, median; box limits, upper and lower quartiles; whiskers, 1.5x interquartile range; points, outliers.

(B) $R^2$ for walking with eyes free (not painted, N=5) and eyes covered (painted, N=5, p-value: 0.34). Box plots show: center line, median; box limits, upper and lower quartiles; whiskers, 1.5x interquartile range; points, outliers.
(C) $R^2$ for walk for all neurons (pan-Gal4;UAS-GCaMP)
(D) $R^2$ for flail for all neurons (pan-Gal4;UAS-GCaMP)
(E) $R^2$ for rest/groom for all neurons (pan-Gal4;UAS-GCaMP)
(F) Correlation coefficient for flailing behavior for all neurons (pan-Gal4;UAS-GCaMP)
(G) Correlation coefficient for grooming behavior for all neurons (pan-Gal4;UAS-GCaMP)
(H) $R^2$ for walking on an air-supported (N=19) vs. a Styrofoam ball (N=13, p-value <0.001). Box plots show: center line, median; box limits, upper and lower quartiles; whiskers, 1.5x interquartile range; points, outliers.
(I) $R^2$ for walking for individual brain regions on an air-supported (N=19) vs. a Styrofoam ball (N=13). Box plots show: center line, median; box limits, upper and lower quartiles; whiskers, 1.5x interquartile range; points, outliers.
(J) Examples of brain activity maps ($R^2$ as z-stacks) and names of individual brain regions
(K) Schematic showing brain regions in adult fly brain.

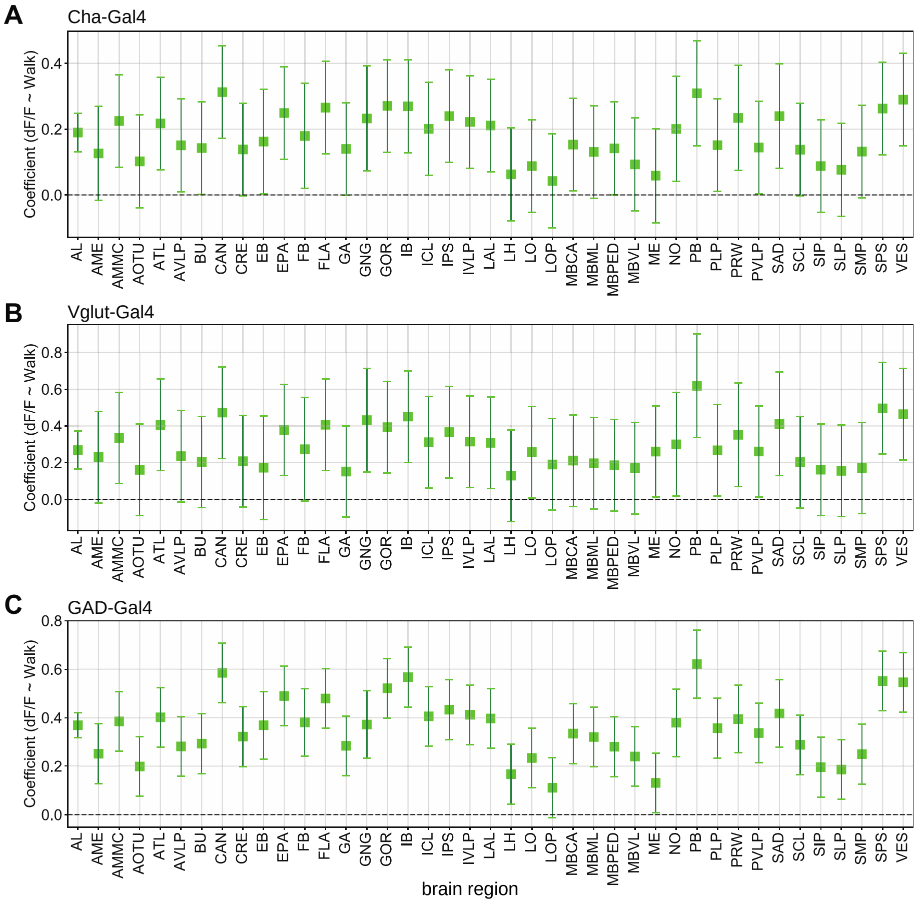


**Fig. S2**
(A) Correlation coefficients during walk for cholinergic neurons (Cha-Gal4;UAS-GCaMP6)
(B) Correlation coefficients during walk for glutamatergic neurons (Vglut-Gal4;UAS-GCaMP6)
(C) Correlation coefficients during walk for GABAergic neurons (GAD-Gal4;UAS-GCaMP6)
N=5 flies for each genotype.

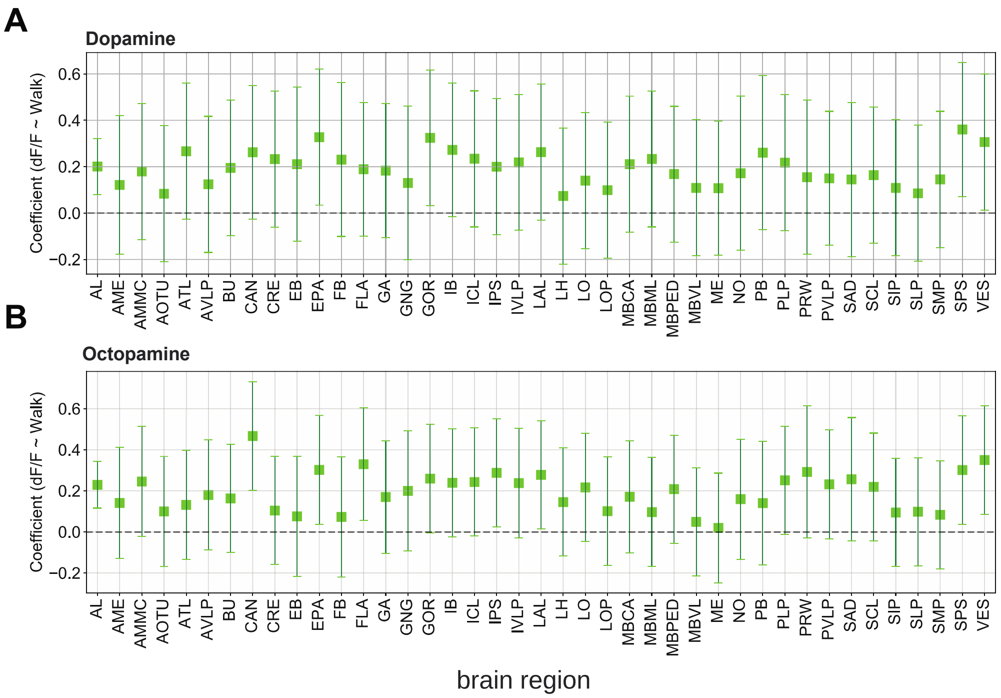


**Fig. S3**

(A) Correlation coefficients during walk for dopaminergic neurons (TH/DDC-Gal4 or GMR58E04-Gal4;UAS-GCaMP6)

(B) Correlation coefficients during walk for octopaminergic neurons (Tdc2-Gal4;UAS-GCaMP6)

TH/DDC-Gal4: N=9, Tdc2-Gal4: N=7, Trh-Gal4: N=6 flies.

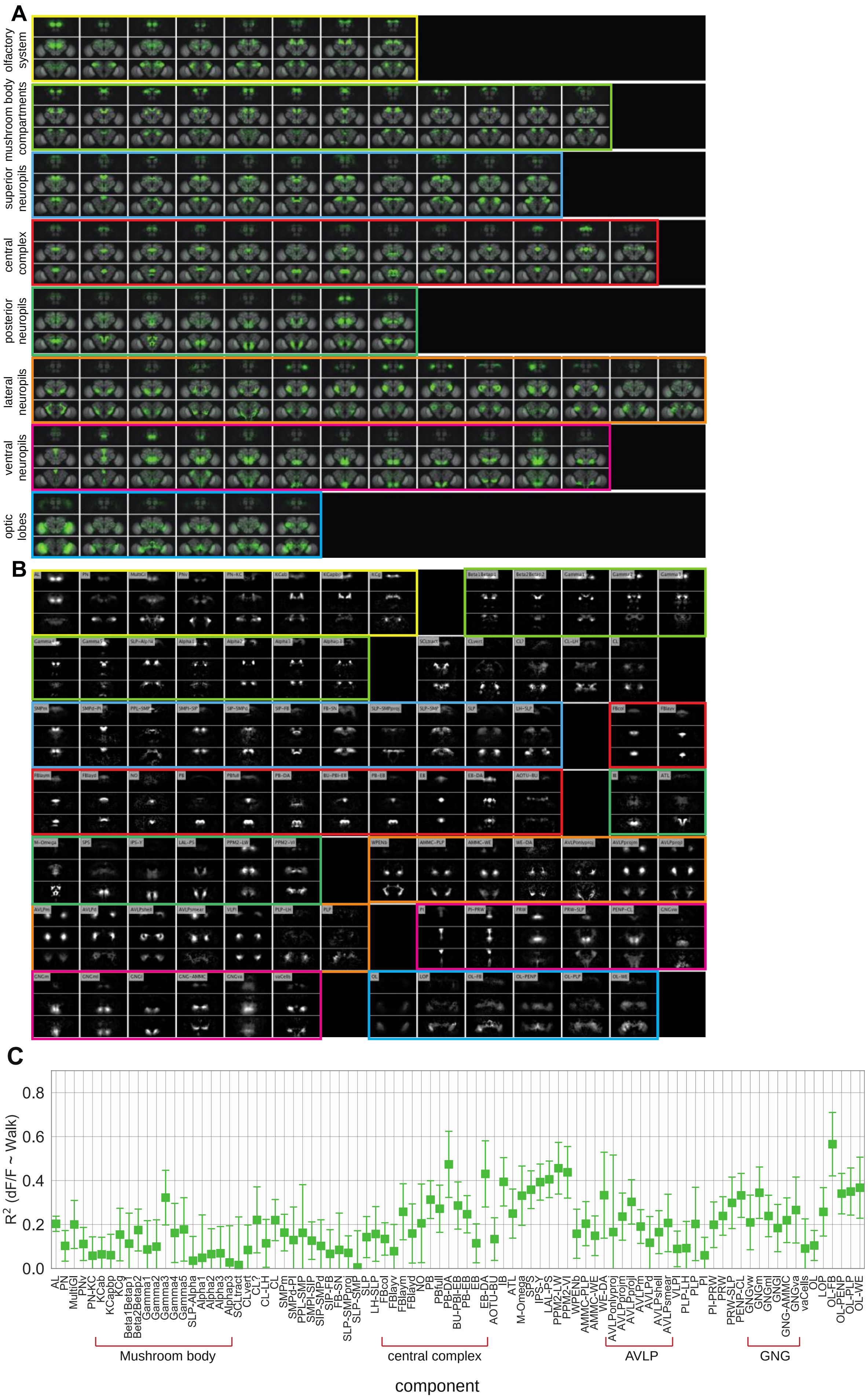

A
olfactory system
mushroom body compartments
superior neuropils
central complex
posterior neuropils
lateral neuropils
ventral neuropils
optic lobes
B
C
0.8
0.6
0.4
0.2
0.0
R² (dF/F ~ Walk)
AL
PN
MultiGl
PNv
PN-KC
KCab
KCapbp
KCg
Beta1Betap1
Beta2Betap2
Gamma1
Gamma2
Gamma3
Gamma4
Gamma5
SLP-Alpha
Alpha1
Alpha2
Alphap3
SCLtract
CLvert
CL?
CL-LH
CL
SMPm
SMPd-PI
PPL-SMP
SMPl-SIP
SIP-SMPd
SIP-FB
FB-SN
SLP-SMPproj
SLP-SMP
SLP
LH-SLP
FBcol
FBlayv
FBlaym
FBlayd
NO
PB
PBfull
PB-DA
BU-PBI-EB
PB-EB
EB
EB-DA
AOTU-BU
IB
ATL
M-Omega
SPS
IPS-Y
LAL-PS
PPM2-LW
PPM2-VI
WPENb
AMMC-PLP
AMMC-WE
WE-DA
AVLPonlyproj
AVLPprojm
AVLPprojl
AVLPm
AVLPd
AVLPshell
AVLPsmear
VLPl
PLP-LH
PLP
PI
PI-PRW
PRW
PRW-SLP
PENP-CL
GNGvw
GNGm
GNGml
GNGl
GNG-AMMC
GNGva
vaCells
OL
LOP
OL-FB
OL-PENP
OL-PLP
OL-WE
Mushroom body
central complex
AVLP
GNG
component

**Fig. S4**

(A) Z-stacks of functional components for walk mapped onto the adult fly brain
(B) Functional components for walk and their names or acronyms
(C) $R^2$ for components activity vs walk. See Table 2 for definition of acronyms. N=58 flies of different genotypes, see table in methods for details.

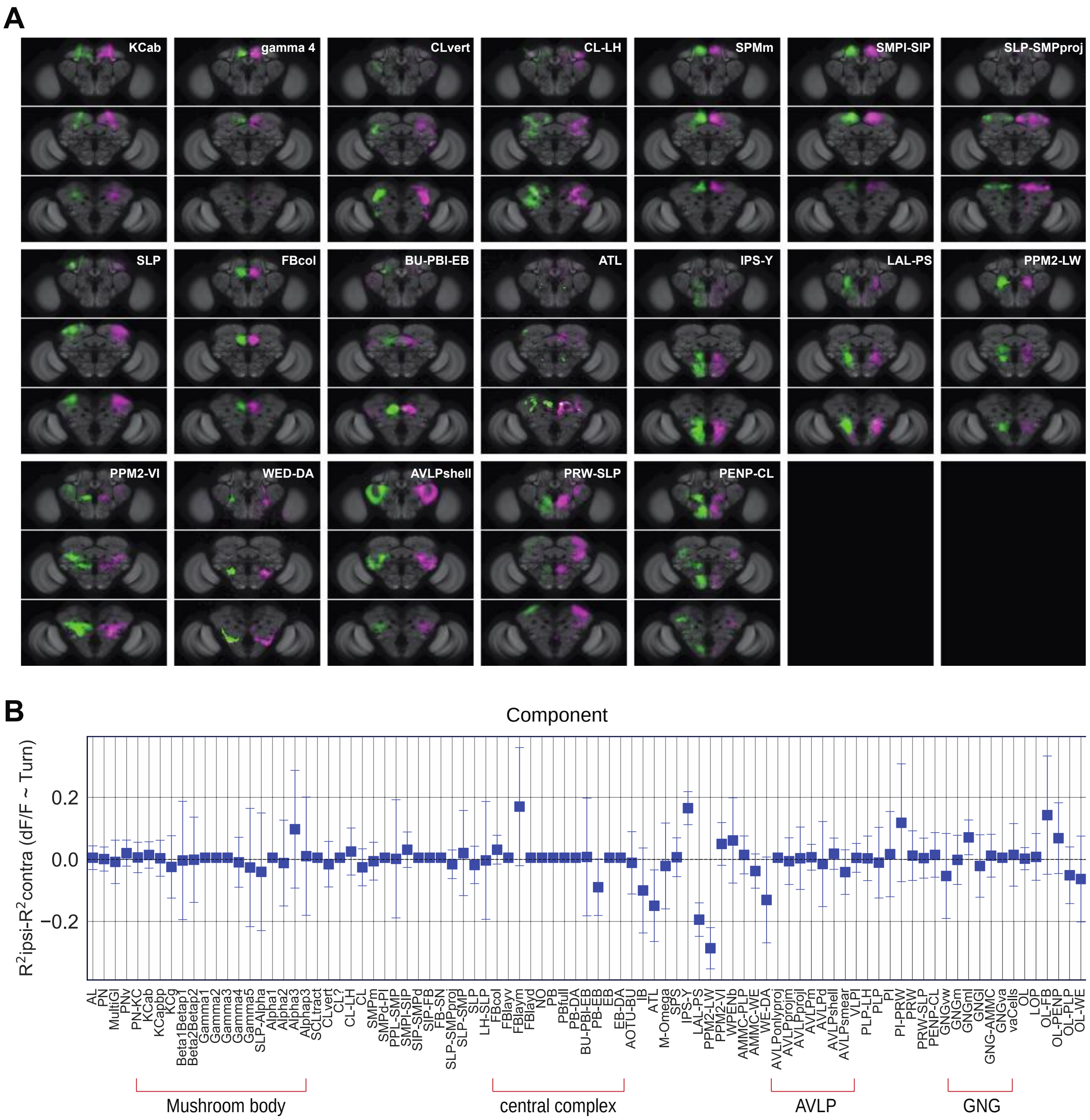


**Fig. S5**

Turning activates specific components and neurons.

(A) Examples of components present in both the left and right hemisphere labeled in different colors (magenta and green).

(B) $R^2$-value of regression coefficients during turn for all genotypes. The value for the contralateral side was subtracted from the ipsilateral side and plotted for every component. N=58 flies of different genotypes, see table in methods for details.

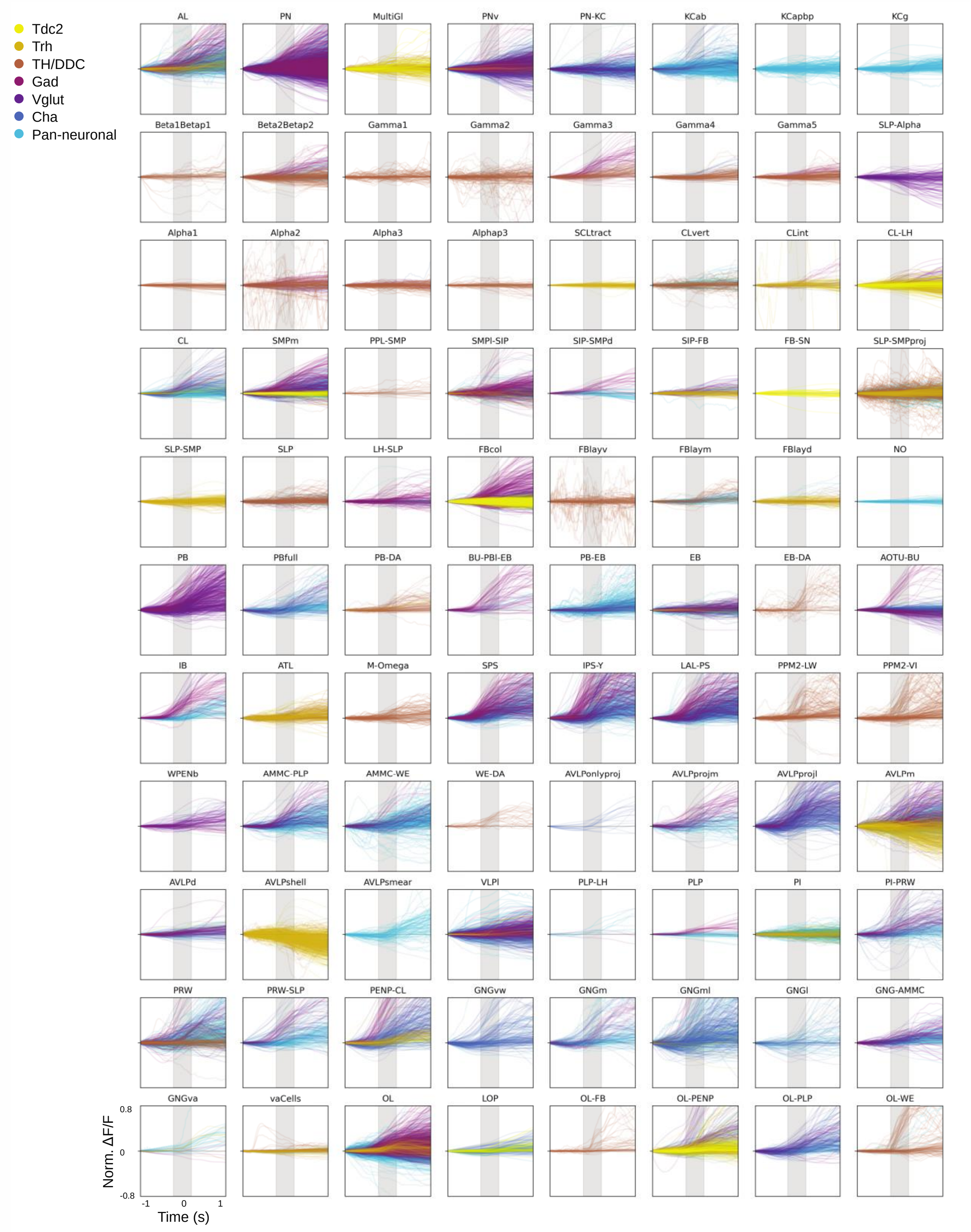
Tdc2
Trh
TH/DDC
Gad
Vglut
Cha
Pan-neuronal
AL
PN
MultiGl
PNv
PN-KC
KCab
KCapbp
KCg
Beta1Betap1
Beta2Betap2
Gamma1
Gamma2
Gamma3
Gamma4
Gamma5
SLP-Alpha
Alpha1
Alpha2
Alpha3
Alphap3
SCLtract
CLvert
CLint
CL-LH
CL
SMPm
PPL-SMP
SMPl-SIP
SIP-SMPd
SIP-FB
FB-SN
SLP-SMPproj
SLP-SMP
SLP
LH-SLP
FBcol
FBlayv
FBlaym
FBlayd
NO
PB
PBfull
PB-DA
BU-PBl-EB
PB-EB
EB
EB-DA
AOTU-BU
IB
ATL
M-Omega
SPS
IPS-Y
LAL-PS
PPM2-LW
PPM2-VI
WPENb
AMMC-PLP
AMMC-WE
WE-DA
AVLPonlyproj
AVLPprojm
AVLPprojl
AVLPm
AVLPd
AVLPshell
AVLPsmear
VLPl
PLP-LH
PLP
Pl
Pl-PRW
PRW
PRW-SLP
PENP-CL
GNGvw
GNGm
GNGml
GNGl
GNG-AMMC
GNGva
vaCells
OL
LOP
OL-FB
OL-PENP
OL-PLP
OL-WE
0.8
0
-0.8
Norm. ΔF/F
-1
0
1
Time (s)

**Fig. S6A**

Graphs show onset of walk trials for all flies and all transgenic lines at all recording speeds. A corrective factor based on regression coefficient with walk (Fluorescence response to walk ~ UAS-GCaMP version: GCaMP7f:0.6,GCaMP6f:1,GCaMP6M:1.1, GCaMP7s 1.2, GCaMP6s:1.4) was applied for direct comparison between experiments with different reporter versions. The average value for the first 50 ms was taken as baseline and subtracted.

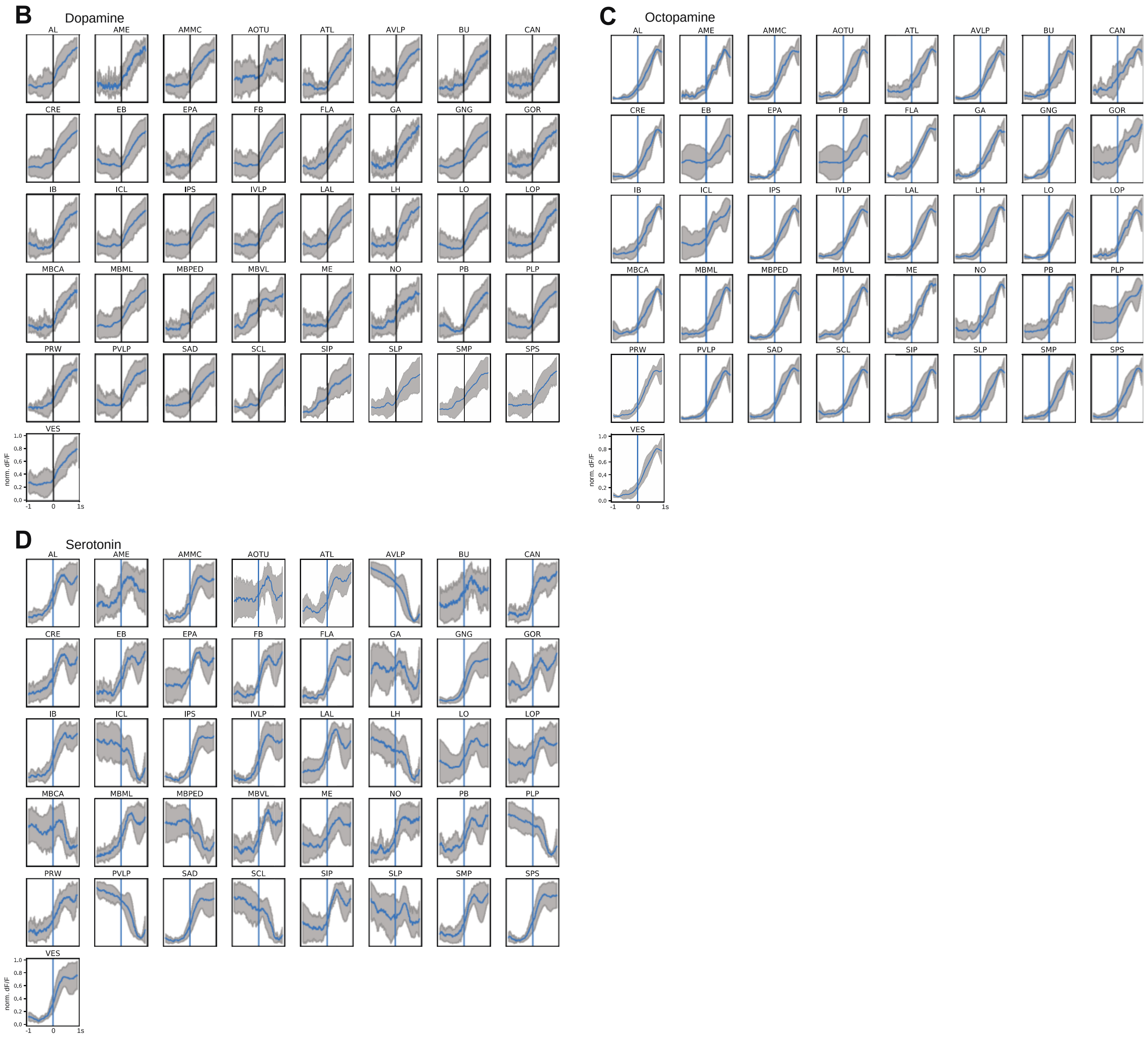


**Fig. S6B-D**

(B) Normalized ΔF/F GCaMP fluorescence for walking-induced activity at 1 s before and 1 s after walk onset (blue line) for different brain regions: For dopaminergic neurons (TH/DDC-Gal4;GCaMP), N=9

(C) For octopaminergic neurons (Tdc2-Gal4;GCaMP), N=7

(D) For serotonergic neurons (Trh-Gal4;GCaMP), N=6 flies.

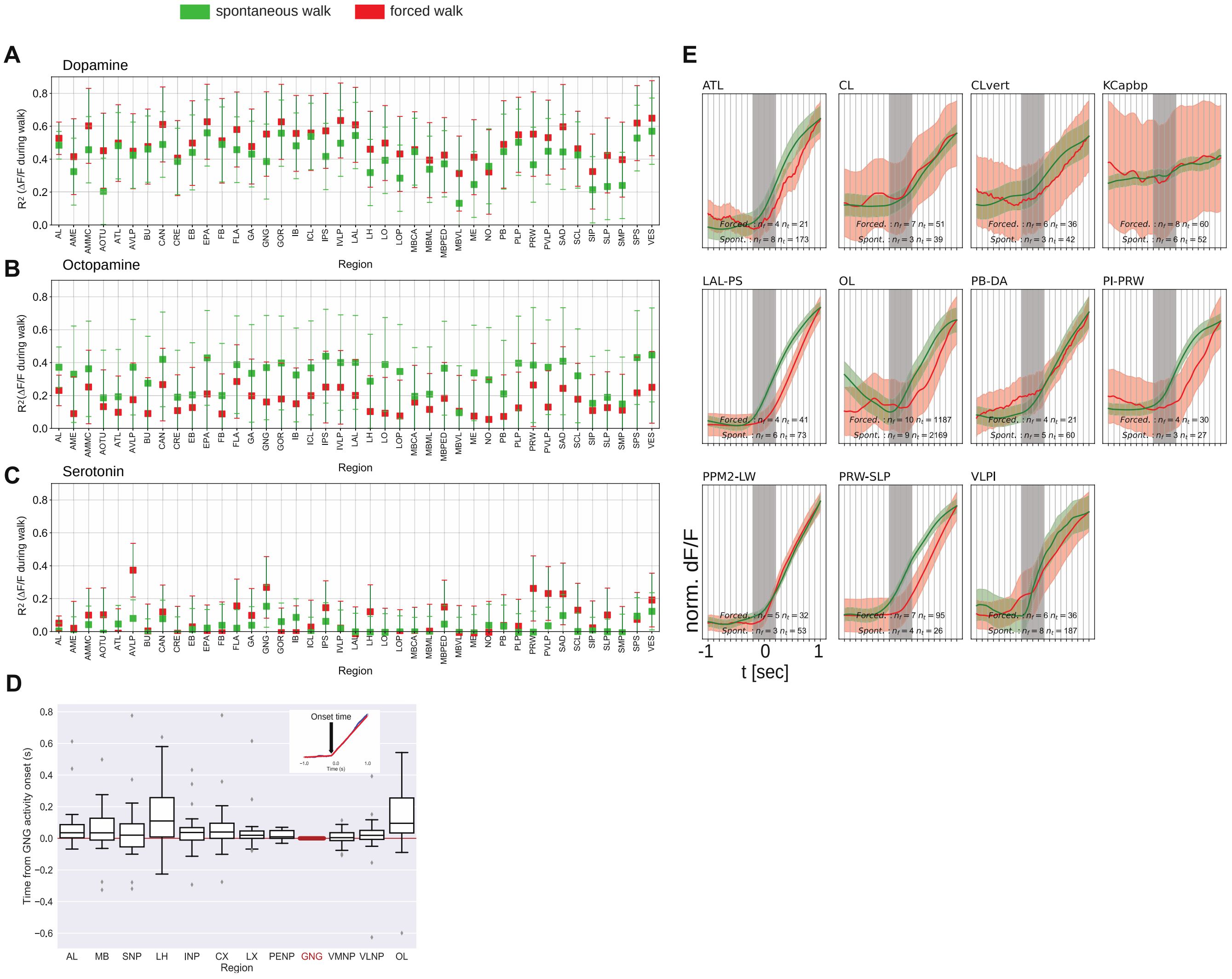


**Fig. S7**

(A) $R^2$ of ΔF/F regression with forced and spontaneous walk for different brain regions for dopaminergic neurons.

(B) $R^2$ of ΔF/F regression with forced and spontaneous walk for different brain regions for octopaminergic neurons.

(C) $R^2$ of ΔF/F regression with forced and spontaneous walk for different brain regions for serotonergic neurons.
TH/DDC-Gal4: spontaneous, N=9, forced, N=5; Tdc2-Gal4: spontaneous, N=7, forced, N=6; Trh-Gal4: spontaneous, N=6, forced, N=6 flies.

(D) Panel showing GCaMP-fluorescence relative to onset of walk for pan-neuronally expressed GCaMPs in different regions relative to the GNG. Piece-wise linear curves were fitted to fluorescence traces 0.5 s before to 0.5 s after the onset of walk. The onset time was determined as the point of change from one linear piece to the next. Note that all medians are positive. Inset: Fit of activity onset. TH/DDC-Gal4: N=9, Tdc2-Gal4: N=7, Trh-Gal4: N=6 flies.

(E) Comparison between activity at walk onset for spontaneous (green) and forced (red) walk, for additional components. Individual trials were normalized and averaged for each component. Shaded regions represent the trail SEM, nf: number of flies, nt: total number of trials.
